# White Matter and Social Cognition

**DOI:** 10.1101/179473

**Authors:** Yin Wang, Athanasia Metoki, Kylie H. Alm, Ingrid R. Olson

## Abstract

There is a growing consensus that social cognition and behavior emerge from interactions across distributed regions of the “social brain”. Social neuroscience has traditionally focused its attention on functional response properties of these gray matter networks and neglected the vital role of white matter (WM) connections in establishing such networks and their functions. In this article, we conduct a comprehensive review of prior research on structural connectivity in social neuroscience and highlight the importance of this literature in clarifying brain mechanisms of social cognition. We pay particular attention to the research on three key social processes: face processing, embodied cognition, and theory of mind, and their respective underlying neural networks. To fully identify and characterize the anatomical architecture of these networks, we further implement probabilistic tractography on a large sample of diffusion-weighted imaging data. The combination of an in-depth literature review and the empirical investigation gives us an unprecedented, well-defined landscape of WM pathways underlying major social brain networks. Finally, we discuss current problems in the field, outline suggestions for best practice in diffusion imaging data collection and analysis, and offer new directions for future research.

**Abbreviations:**

ACC: anterior cingulate cortex
AD: axial diffusivity
AF: arcuate fasciculus
AI: anterior insula
ALS: amyotrophic lateral sclerosis
AMG: amygdala
ASD: autism spectrum disorders
ATL: anterior temporal lobe
ATR: anterior thalamic radiation
CC: corpus callosum
CING: cingulum bundle
CST: cortico-spinal tract
DES: direct electrical stimulation
dMPFC: dorsal medial prefrontal cortex
dMRI: diffusion-weighted MRI
DP: developmental prosopagnosia
DTI: diffusion tensor imaging
FA: fractional anisotropy
FFA: fusiform face area
IFG: inferior frontal gyrus
IFOF: inferior fronto-occipital fasciculus
ILF: inferior longitudinal fasciculus
IPL: inferior parietal lobe
MCI: mild cognitive impairment
MD: mean diffusivity
MPFC: medial prefrontal cortex
MS: multiple sclerosis
OFA: occipital face area
OFC: orbitofrontal cortex face patch
PCC: posterior cingulate cortex
PD: Parkinson’s disease
PP: progressive prosopagnosia
PreC: precuneus
RD: radial diffusivity
ROI: region-of-interest
sMRI: structural MRI
STS: superior temporal sulcus
TBSS: tract-based spatial statistics
ToM: Theory of Mind
TPJ: temporo-parietal junction
UF: uncinate fasciculus
VBM: voxel based morphometry
vMPFC: ventral medial prefrontal cortex
WM: white matter

## Introduction

Just as H.M. arguably became the index case for the neuroscience of memory, so may Phineas Gage serve as the index case for the neuroscience of social cognition (Kihlstrom, 2010). The notable case of Phineas Gage appears in every textbook of social neuroscience, and illustrates the critical role of the frontal cortex in human personality and social behavior. However, a recent study (Van Horn et al., 2012) reconstructed Gage’s injury and revealed a new picture of his brain damage: while 4% of the frontal cortex was intersected by the rod’s passage, 11% of frontal white matter (WM) was also damaged. Fasciculi that were severely impacted include the uncinate fasciculus (UF), cingulum bundle (CING), and superior longitudinal fasciculus (SLF), and they likely played crucial roles in Gage’s reported post-injury social and behavioral changes. Clearly, the significance of WM for social cognition has been profoundly underestimated since the beginning of social neuroscience.

### Aim of the Article

The past few years have seen an increasing number of structural connectivity studies of social cognition, many of them propelled by the fast development of diffusion imaging techniques. Despite this, no dedicated review or meta-analysis exists in this field. The purpose of this review article is five-fold. First, we present opinions on why social neuroscience needs research on WM connectivity. Second, we provide a brief introduction to the approaches and methods used in this research. Next, we provide an overview of the field by surveying 51 existing studies of WM related to three key social processes (face processing, embodied cognition, and theory of mind) and their respective underlying brain networks (face perception network, mirroring network, and mentalizing network). Detailed information about these studies, such as sample size, experimental protocols, analysis methods, and associated WM tracts are summarized and discussed. In addition, to better understand the WM connectivity profile within each network, we further carry out an empirical investigation by using probabilistic tractography on a large diffusion-weighted imaging dataset. We then make conclusions based on the convergence of findings across the survey and the empirical study. Finally, we outline current problems in the field, discuss emerging trends in methodology, and highlight new directions for future research. To our knowledge, this is the only review to examine the influence of WM in social neuroscience, as well as the most comprehensive empirical study to-date to elucidate the connectivity profile of social brain networks.

### Why Do We Study WM in Social Neuroscience?

The history of human neuroscience shows an overwhelming emphasis on the functionality of gray matter, with a relative disregard of WM (Figure 1). Students are taught to identify sulcal-gyral landmarks that denote gray matter regions; however, the identification of WM tracts is ignored. Similarly, studies of individuals with focal brain lesions due to stroke or resection often show involvement of WM in brain scans, but rarely is WM damage an explicit focus of the discussion, and in some cases, its involvement in the cognitive process being examined is discounted. However, few would deny the importance of WM for human cognition and behavior. It makes up half of the whole cerebral volume and plays a vital role in communications between cortical areas (Douglas Fields, 2008; Schoenemann, Sheehan, & Glotzer, 2005). Studies of human WM can provide insight into the organization of brain systems and the functions they perform. Several WM structures (e.g. optic tract, corticospinal tract, fornix, arcuate fasciculus) have been well characterized for vision, motor skills, memory, attention, and language (Doricchi, Thiebaut de Schotten, Tomaiuolo, & Bartolomeo, 2008; Fields, 2008; Filley & Fields, 2016; Friederici, 2015; Kleinschmidt & Vuilleumier, 2013; Rokem, Bock, Scherf, Wandell, & Bridge, 2017; Thomas, Koumellis, & Dineen, 2011). However, current knowledge about the specific WM tracts underlying social cognition is limited.

**Figure 1.**
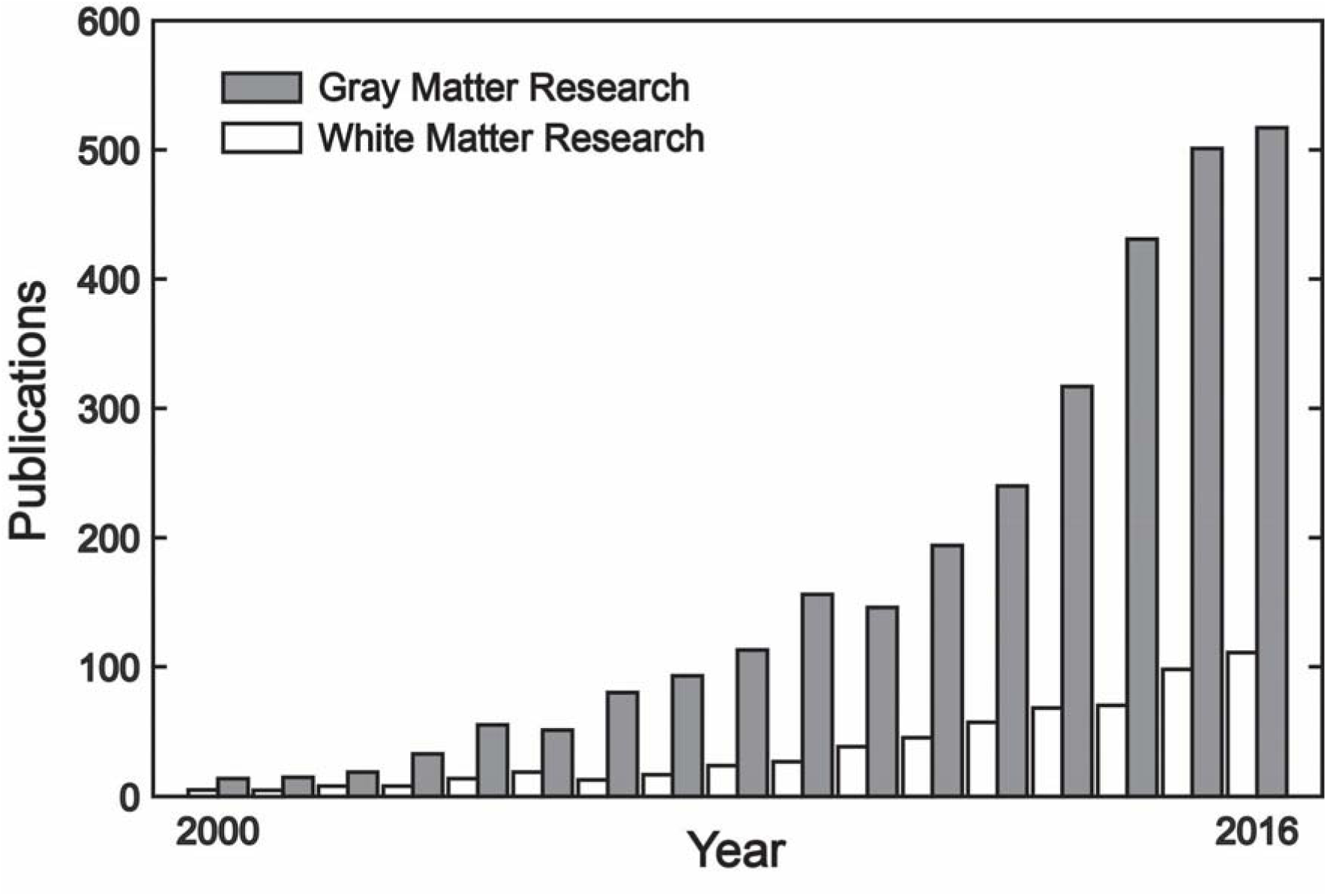
The proliferation of gray matter and white matter studies in social neuroscience. Both types of research have been rapidly increased over the past 15 year; however, the number of white matter studies per year is always less than 1/3 of the number of gray matter studies. The plotted data were extracted from https://www.ncbi.nlm.nih.gov/pubmed/ on 4/30/2017, using the search term “(social) *AND* (gray matter *OR* fMRI *OR* functional imaging)” for gray matter research (gray bars) and “(social) *AND* (white matter *OR* DTI *OR* diffusion imaging)” for white matter research (white bars).

This issue becomes even more pressing as the field of social neuroscience is increasingly moving away from a strict modular view of the brain and toward an appreciation of circuits and networks (i.e. “connectionism”; Bassett & Sporns, 2017). Although early research attempted to locate social cognitive processes in individual brain regions, such as medial prefrontal cortex and temporo-parietal junction, there is a growing consensus that social processes and behavior emerge from interactions across distributed areas of the so-called “social brain networks” (Kennedy & Adolphs, 2012). Correspondingly, this network-based approach features a variety of functional connectivity analyses (e.g. psychophysiological interaction, resting-state, dynamic causal modeling) examining the functional interactions between social brain areas. Yet, these methods are solely based on gray matter co-activation (i.e. BOLD signal). However, the presence of WM connectivity is a prerequisite for the existence of functional networks and ensuing rapid, efficient interactions (Jbabdi, Sotiropoulos, Haber, Van Essen, & Behrens, 2015), and the properties of anatomical connectivity, to some degree, determine the properties of functional connectivity (Boorman, O’Shea, Sebastian, Rushworth, & Johansen-Berg, 2007; Bullmore & Sporns, 2009; Honey et al., 2009). Some social disorders (e.g. developmental prosopagnosia) appear to be mainly caused by damage to WM tracts between social brain areas, while functional activation within gray matter regions remains relatively intact (Avidan et al., 2014; Behrmann, Avidan, Marotta, & Kimchi, 2005; Thomas et al., 2009). Therefore, our understanding of social cognition and social dysfunction will be incomplete until we understand the anatomical connectivity of the social brain. This is precisely why recent large-scale coordinated brain projects, such as the Human Connectome Project (Glasser et al., 2016; Van Essen et al., 2013) and UK biobank (Miller et al., 2016), incorporate rigorous measurement of structural connectivity in their pipelines.

Complementary to functional approaches (e.g. fMRI, EEG, MEG, TMS), WM investigation provides crucial insights into the neural basis of social cognition. First, *in vivo* imaging of WM bundles allows us to unveil the structural architecture of the social brain. Diffusion-weighted MRI tractography methods make it possible to virtually dissect the social brain and explore patterns of WM projections between social brain regions (Gschwind, Pourtois, Schwartz, Van De Ville, & Vuilleumier, 2012; Pyles, Verstynen, Schneider, & Tarr, 2013). When combined with functional MRI localizers, tractography can delicately trace the specific WM pathway underlying communications between functional regions of a particular social process, such as gaze perception (Ethofer, Gschwind, & Vuilleumier, 2011) and imitation (Hamzei et al., 2016).

Second, human social behavior displays remarkable variance between individuals. Research focused on variation in WM opens up new opportunities to explore the structural basis of such inter-individual variability (Johansen-Berg, 2010; Kanai & Rees, 2011). WM properties have been successfully used to account for diversity in personality traits (Cohen, Jan-Christoph, Schoene-Bake, Elger, & Weber, 2008; Iidaka, Miyakoshi, Harada, & Nakai, 2012; Kim & Whalen, 2009; Lei et al., 2014; Xu & Potenza, 2012), self-processes (Chester, Lynam, Powell, & DeWall, 2016; Wang et al., 2014; Yang et al., 2016) and social network size (Hampton, Unger, Von Der Heide, & Olson, 2016; Molesworth, Sheu, Cohen, Gianaros, & Verstynen, 2014), and to predict social skills, such as face recognition ability (Tavor et al., 2014), social memory (Metoki, Alm, Wang, Ngo, & Olson, 2017; Unger, Alm, Collins, O’Leary, & Olson, 2016) and empathy (Chou, Cheng, Chen, Lin, & Chu, 2011; Parkinson & Wheatley, 2014; Takeuchi et al., 2013). Associations between social impairments in various clinical patients and WM indices sometimes make even better predictions than regional gray matter indices (Downey et al., 2015).

Third, research on WM connectivity that relates brain to behavior plays a key role in unraveling the structural correlates of social behavior changes during development (Anderson, Rice, Chrabaszcz, & Redcay, 2015; Chavez & Heatherton, 2017; Grosse Wiesmann, Schreiber, Singer, Steinbeis, & Friederici, 2017; Scherf, Thomas, Doyle, & Behrmann, 2014), aging (Cabinio et al., 2015; Charlton, Barrick, Markus, & Morris, 2009), disease, (Baggio et al., 2012; Philippi, Mehta, Grabowski, Adolphs, & Rudrauf, 2009) and training/intervention processes (Thomas & Baker, 2013). It has been consistently found that maturation or deterioration of WM throughout the lifespan relates to development or decline in social skills (Johansen-Berg, 2010; Scherf et al., 2014; Thomas et al., 2008). In addition, WM changes can be associated with phylogeny of social functions. For example, cross-species comparison of WM pathways has helped to reveal the anatomical signature of certain evolved social skills, such as imitation and theory of mind (Hecht et al., 2013; Mars, Neubert, et al., 2012; Rilling et al., 2008). It is worth mentioning that WM imaging techniques (i.e. diffusion and structural MRI) are easy to carry out for special groups who may not tolerate long scan times or challenging tasks (e.g. children, seniors, patients, and non-human animals).

Furthermore, our understanding of certain psychiatric and neurological disorders can significantly benefit from WM research in social neuroscience. Multiple psychiatric and neurological disorders feature a primary impairment in social cognition (e.g. autism and developmental prosopagnosia) and many others also show abnormalities in social functioning (e.g. schizophrenia). As already mentioned above, the network-based approach prompts clinical researchers to consider connectivity rather than focal damage when interpreting social dysfunction, and the health of WM connections is an important factor underlying many social disorders (Kennedy & Adolphs, 2012). Since WM microstructure and connectivity in healthy populations can be used as a baseline for comparison with clinical populations with social deficits, these structural patterns can be treated as promising biomarkers for clinical diagnosis and therapy (Dopper et al., 2013; Johansen-Berg, 2010; Woo, Chang, Lindquist, & Wager, 2017). By linking anatomical WM characteristics to clinical phenotypes (e.g. neuropsychological standardized tests), one can investigate the impact of brain pathology on social behavior via WM disruptions (Jalbrzikowski et al., 2014; Levin et al., 2011; Mike et al., 2013; Scheibel et al., 2011) or study the etiology of social disorders (Ameis & Catani, 2015; Thomas et al., 2009; Travers et al., 2012). With the ongoing development of diffusion and quantitative MRI (Alexander et al., 2011; Wandell, 2016), more subtle WM properties and changes can be detected in each disorder, making the WM approach particularly fruitful for studying network-level anatomy, functioning, plasticity, and compensation in the social brain (Kennedy & Adolphs, 2012).

Finally, studying WM enables us to infer “structure-function” relationships that subserve social cognition and behavior. For face processing, structural maturation of WM tracts during early development is necessary for functional specialization of the face network to emerge. It has been found that increasing WM integrity in face pathways (via myelination) increases the propagation of the neural signal throughout the face perception network, thereby enhancing the functional characteristics of the nodes within the network (Scherf et al., 2014). In addition, the function that a social brain region serves is often dynamic, and changes across networks (Yang, Rosenblau, Keifer, & Pelphrey, 2015); even for the same region, different subareas might belong to different anatomical networks and thus play different functional roles (Mars, Sallet, et al., 2012). By using WM connectivity profiles, we are now able to identify boundaries of functionally distinct brain areas (i.e. tractography-based parcellation). This “connectional localizer” approach (Jbabdi et al., 2015) has been successfully employed in functional segmentation of multiple social brain areas, including medial prefrontal cortex (Johansen-Berg et al., 2004; Sallet et al., 2013), temporo-parietal junction (Mars, Sallet, et al., 2012), inferior frontal gyrus (Tomassini et al., 2007), superior temporal sulcus (Xu et al., 2016), inferior parietal lobe (Mars et al., 2011), insula (Ghaziri et al., 2017) and occipital face area (Pyles et al., 2013). When combined with powerful machine learning algorithms, WM connectivity fingerprints can even predict idiosyncratic functional responses to social stimuli (Saygin et al., 2011).

### Social Cognition and Brain Networks

Social cognition is the collection of mental processes that allow individuals to interact with one another (Frith & Frith, 2007, 2012). An extensive literature in social neuroscience suggests that there are at least three large-scale neural networks/circuits underlying social processes and interactions (Cross, Ramsey, Liepelt, Prinz, & Hamilton, 2016; Kennedy & Adolphs, 2012; Van Overwalle, 2009; Van Overwalle & Baetens, 2009; Yang et al., 2015): the “face perception network” (Duchaine & Yovel, 2015; Gobbini & Haxby, 2007; Haxby, Hoffman, & Gobbini, 2000), the “mirroring network” (Bonini, 2017; Iacoboni, 2009a; Molenberghs, Cunnington, & Mattingley, 2012; Rizzolatti & Craighero, 2004) and the “mentalizing network” (Mar, 2011; Schurz, Radua, Aichhorn, Richlan, & Perner, 2014) (see Figure 2).

**Figure 2.**
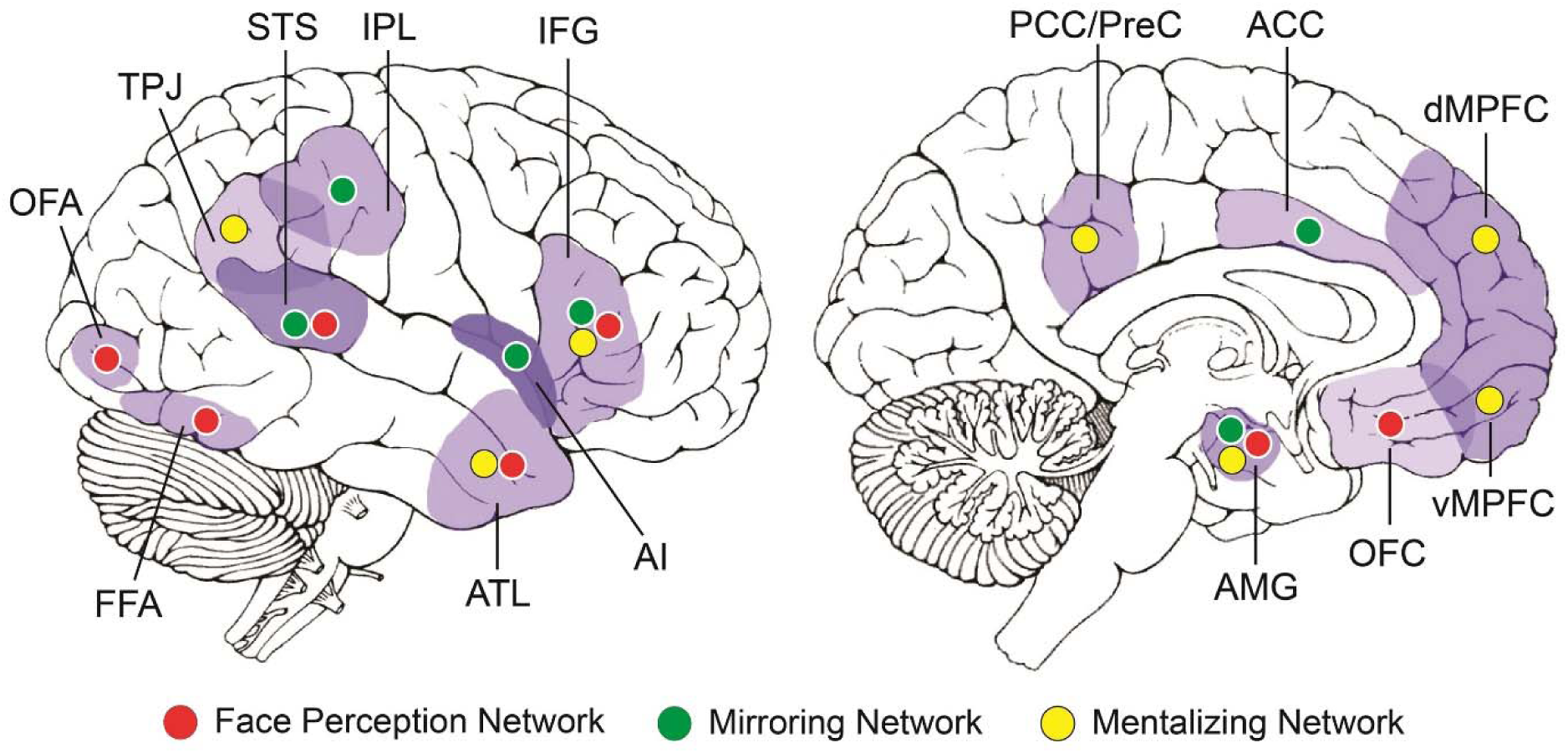
Three major networks in the social brain. ACC, anterior cingulate cortex; AI, anterior insula; AMG, amygdala; ATL, anterior temporal lobe; dMPFC, dorsomedial prefrontal cortex; FFA, fusiform face area; IFG, inferior frontal gyrus; IPL, inferior parietal lobule; OFC, orbitofrontal cortex; OFA, occipital face area; PCC/PreC, posterior cingulate cortex/precuneus; STS, superior temporal sulcus; TPJ, temporoparietal junction; vMPFC, ventromedial prefrontal cortex.

### Face Processing and Face Perception Network

Social interactions often start with recognizing conspecific’s faces. This ability is arguably the most developed social skill in humans. Converging empirical evidence suggests that face perception is mediated by a widely distributed network of face-selective areas, each engaging in different aspects of face processing (Duchaine & Yovel, 2015; Gobbini & Haxby, 2007; Haxby et al., 2000; Ishai, Schmidt, & Boesiger, 2005). For example, posterior regions, such as the occipital face area (OFA), process low-level visual features and analyze facial parts (Pitcher, Walsh, & Duchaine, 2011); the fusiform face area (FFA) is involved in processing invariant facial features, such as identity and gender (Haxby et al., 2000), whereas the posterior superior temporal sulcus (STS) is more sensitive to changeable features, such as facial expression and lip movement (Gobbini & Haxby, 2007). Anterior regions, such as the amygdala (AMG), subserve emotional aspects of face representations (Mende-siedlecki, Said, & Todorov, 2013); the anterior temporal lobe (ATL) stores conceptual knowledge related to faces, including names and biographical information (Collins & Olson, 2014; Wang et al., 2017); the inferior frontal gyrus (IFG) processes the semantic aspects of faces as well as gaze directions (Chan & Downing, 2011; Duchaine & Yovel, 2015; Ishai, 2008), and the orbitofrontal cortex (OFC) evaluates rewarding aspects of faces, like facial attractiveness and trustworthiness (Mende-siedlecki et al., 2013; Troiani, Dougherty, Michael, & Olson, 2016).

### Embodied Cognition and Mirroring Network

Social interactions also require individuals to rapidly and effortlessly grasp others’ intentions and emotions, and respond accordingly and appropriately. These social reciprocity skills are often linked to the so-called “mirroring network”, which mediates our capacity to share the meaning of actions and emotions through the embodied simulation mechanism (Gallese, 2007). By simulating observed action (or emotions) with one’s own motor (or affective) system, the mirroring mechanism provides the basis for action understanding (Rizzolatti, Cattaneo, Fabbri-Destro, & Rozzi, 2014; Rizzolatti & Craighero, 2004; Rizzolatti & Sinigaglia, 2010), imitation (Caspers, Zilles, Laird, & Eickhoff, 2010; Iacoboni, 2009b), emotional recognition (Banissy et al., 2011; Bastiaansen, Thioux, & Keysers, 2009; Braadbaart, de Grauw, Perrett, Waiter, & Williams, 2014; Niedenthal, Mermillod, Maringer, & Hess, 2010; van der Gaag, Minderaa, & Keysers, 2007; Wood, Rychlowska, Korb, & Niedenthal, 2016) and empathy (Bernhardt & Singer, 2012; Corradini & Antonietti, 2013; Gonzalez-Liencres, Shamay-Tsoory, & Brune, 2013; Iacoboni, 2009a; Shamay-Tsoory, 2011). In humans, the putative mirroring network is formed by a collection of areas (Iacoboni, 2009a; Molenberghs et al., 2012; Mukamel, Ekstrom, Kaplan, Iacoboni, & Fried, 2010; Rizzolatti & Craighero, 2004), including the inferior frontal gyrus (IFG, which represents motor plans of actions; Rizzolatti et al., 2014), the inferior parietal lobule (IPL, which represents abstract action goal; Hamilton & Grafton, 2006), the posterior STS (which is theorized to serve as the sensory input of the network; Rizzolatti & Craighero, 2004), the anterior cingulate cortex (ACC, empathy for pain; Bernhardt & Singer, 2012), the anterior insula (AI, empathy for disgust; Bernhardt & Singer, 2012) and the amygdala (AMG, empathy for fear; Bastiaansen et al., 2009).

### Theory of Mind and Mentalizing Network

Finally, the capacity to make accurate inferences about the mental states of other people (e.g. their thoughts, needs, desires, and beliefs) is important for predicting the behavior of others and for facilitating social interactions (Blakemore, 2008). This particular skill and its associated mental processes have often been referred to as “mentalizing” or “theory of mind” (ToM). A large number of neuroimaging and lesion studies have delineated an extensive brain network for mentalizing abilities (Figure 2), mainly including the dorsal and ventral medial prefrontal cortex (dMPFC and vMPFC), the temporo-parietal junction (TPJ), the posterior cingulate cortex/precuneus (PCC/PreC), the ATL, the IFG and the AMG (Mar, 2011; Molenberghs, Johnson, Henry, & Mattingley, 2016; Schurz et al., 2014; Van Overwalle, 2009; Van Overwalle & Baetens, 2009). The specific function of each region has not yet been clarified, but some (e.g. MPFC and TPJ) are consistently engaged irrespective of the mental state contents and the task modalities (Schurz et al., 2014), whereas the involvement of other regions seems to be more task-dependent (Carrington & Bailey, 2009; Molenberghs et al., 2016).

### How Do We Study WM?

Before stepping into the literature review, we provide a brief introduction to measurements and methods that researchers typically employ to study human WM. As a misconception, it had often been implicitly assumed that WM connections have a binary nature like electrical cables, either they are connected and functioning or they are disconnected. We now know that it is much more nuanced with macrostructural (e.g. fiber’s size, shape and density) and microstructural properties (e.g. fiber’s integrity and degree of myelination) contributing to the functional profile of WM connections.

A wealth of neuroanatomical tools has been developed to measure these macro- and microstructural properties. Tracing techniques are traditionally considered to be the gold standard in measuring WM connections. They can give us extremely detailed, precise connectivity information with low false positive rates. However, they are not ideal for exploring WM connections underlying human social cognition, because (1) they are invasive tools that are restricted to postmortem and animal research; (2) they can only measure a very small percentage of brain connections (let along the large-scale social brain networks); and (3) they cannot be used for multimodal investigations (e.g. in conjunction with functional or behavior measurements, or longitudinal analyses) (Jbabdi et al., 2015). In contrast, albeit far less precise, non-invasive techniques using magnetic resonance imaging (MRI) allow us to measure brain-wide long-range WM connections *in vivo*, in addition to the potential for parallel acquisition of other types of data that can be related to or predicted by WM connectivity.

In general, there are three major tools that have been used to measure WM connections in social neuroscience: diffusion-weighted MRI (dMRI), structural MRI (sMRI), and direct electrical stimulation (DES). In principle, dMRI is mainly used for characterizing macro- and microstructural properties of WM tracts, as well as for delineating long-range WM pathways between disparate brain regions. sMRI makes it possible to visualize and evaluate the macroscopic properties of local WM at high resolution, which is ideal for anatomical morphometry and detection of WM abnormalities and damage for clinical diagnosis. DES provides real-time causality investigations on the functional role of WM tracts.

### Diffusion-Weighted MRI (dMRI)

dMRI is the most popular and powerful technique for exploring WM anatomy and quantifying WM properties in the living human brain. The basic principle and concept behind this technique is that dMRI measures the random motion or diffusion of water molecules, which is restricted by tissue microstructure. When this microstructure is more organized, such as in WM, water diffusion is anisotropic, in that diffusion is less hindered parallel than perpendicular to WM fibers. Thus, by measuring the orientational dependence of water diffusion, dMRI infers the microstructure and properties of surrounding WM tissue (Jbabdi et al., 2015).

The simplest way to quantify the degree of anisotropic diffusion is the diffusion tensor model, which estimates the diffusion process by an ellipsoid, also known as tensor (hence the name origin of diffusion tensor imaging, DTI) (Soares, Marques, Alves, & Sousa, 2013). This model has been so influential that DTI is often used as a synonym for dMRI. Several metrics can be derived from DTI in each voxel, including the mean diffusivity (MD), the degree of anisotropy (i.e. factional anisotropy, FA) and two directional diffusivity measures (i.e. axial diffusivity, AD; radial diffusivity, RD). Variations in these metrics have been associated with alterations in the underlying WM microstructure. While FA is often used as the primarily measure of WM connectivity strength and integrity, MD/AD/RD are useful indicators of WM maturation and dysfunction (e.g. demyelination and axonal damage) (Alexander, Lee, Lazar, & Field, 2007). Four analysis strategies are typically applied to DTI metrics when researchers try to identify local WM differences across individuals or abnormalities in clinical populations (Soares et al., 2013): they can be compared locally in every voxel after registration to an anatomical atlas (voxel-based analysis), or averaged within *a priori* specific regions of interest (ROI-based analysis), or sampled along pathways after fiber tract reconstruction (tractography-based analysis), or analyzed based on the skeletonization of group registered FA maps (tract-based spatial statistics, TBSS) (Feldman, Yeatman, Lee, Barde, & Gaman-Bean, 2010; Soares et al., 2013; Travers et al., 2012). These strategies can also be applied to the investigation of anatomical correlates of numerous experimental and clinical conditions. An in-depth exploration of the relative strengths and weaknesses of each method can be found in several review papers (Feldman et al., 2010; Jones & Cercignani, 2010; Soares et al., 2013).

Another advantage of dMRI techniques is the ability to visualize and characterize long-range WM pathways. To date, dMRI tractography is the only available tool to estimate the trajectories of WM fibers *in vivo*, by measuring the principal direction of water diffusivity on a voxel-by-voxel basis and piecing together information from contiguous voxels (Jbabdi et al., 2015). There are two types of tractography algorithms: deterministic and probabilistic. Deterministic tractography is designed to trace a single path between two regions of interest, and thus is more suitable for identifying large WM fasciculi of the brain. Probabilistic tractography is more useful for quantitatively analyzing the connectivity between two regions based on the probability of a connection, taking into account that a single voxel might connect with more than one target voxel (Roberts, Anderson, & Husain, 2013; Rokem et al., 2017). Once dMRI tractography is completed for a particular WM pathway, one can inspect its macroscopic features (e.g. trajectory shape and volume), microstructural properties (e.g. FA/MD/AD/RD) and connectivity strength (e.g. probability or streamline count) (Soares et al., 2013). These approaches allow the researcher to compare equivalent WM pathways across individuals, even if the precise location of the tract varies (Feldman et al., 2010).

A fundamental limitation of dMRI is the indirect nature of its measurements. Since all estimates are based on water diffusivity, dMRI techniques provide only computational models of WM tissues with many theoretical assumptions about the underlying processes and structures. This makes dMRI error-prone and highly dependent on the data quality, the chosen diffusion model, and the analysis pipeline used (Jones, Knosche, & Turner, 2013).

### Structural MRI (sMRI)

Conventional MRI techniques can also provide useful qualitative and quantitative measurements of WM structures in the brain. Due to its availability and ease of use, sMRI is widely used in clinical research, particularly for WM pathologies (e.g. multiple sclerosis and diffuse low-grade glioma). Rather than measuring water diffusion rate, sMRI collects MR signals (T1 or T2 relaxation) that vary across tissue types, since gray matter contains more cell bodies while WM is primarily composed of myelinated axons and glial cells. sMRI with morphometric analysis is used to measure the shape, size, myelination, and integrity of WM structures, which is very helpful for quantitative assessments of local WM changes in patient-control studies. Additionally, further combination with lesion-symptom mapping can reveal the anatomical basis of abnormal behavior. One limitation of sMRI is that this technique only allows for voxel-level analysis, which restricts investigations to local WM characteristics. In addition, sMRI alone primarily measures the macroscopic morphology of WM tissues, with little information about microstructural properties (unless using very sophisticated modeling such as multi-compartment models) (Jbabdi et al., 2015). A recent trend is to complement sMRI with dMRI. By using both macrostructural measurements from sMRI (e.g. WM volume) and microstructural metrics from dMRI (e.g. FA), one can get a comprehensive picture of WM maturation and integrity (Erus et al., 2015).

### Direct Electrical Stimulation (DES)

dMRI and sMRI primarily use correlation analyses to reveal the relationship between WM tracts and behavior. Correlation is clearly not causation, and further investigations, such as DES, are needed to validate the structure-function relationship. The use of DES on patients during awake neurosurgery provides a unique opportunity to gain insight into the function of WM tracts implicated in human cognition (Duffau, 2015). Electrical stimulation to a spatially and topographically well-defined WM tract creates a “virtual” lesion, which disrupts the function of that WM tract and consequently changes corresponding behavior. This technique provides real-time structure-function mapping with high spatial resolution, and has great advantages in scrutinizing the exact role (i.e. critical versus participatory) of a particular WM tract for a specific mental process.

Like other techniques, DES has several inherent problems. First, because the technique is invasive (e.g. partial resection is required to access the WM beyond the cortex) and only restricted to special clinical groups (e.g. patients with gliomas), the sample sizes in DES studies are typically small. Second, some patients (e.g. with low-grade glioma) may have exhibited abnormal WM profiles for a prolonged period of time, thus confounding DES results with neuroplasticity and compensation effects. Third, the range of behavioral assessment is often limited in DES research, due to limited time available during surgery (Duffau, 2015).

## A Literature Review of WM Research in Social Neuroscience

### Literature Search

So far, no systematic review or meta-analysis has been conducted on the topic of social cognition and WM. In order to get an overview of this body of work, particularly those studies related to the three major social brain networks, we performed a comprehensive online literature search on PubMed, Web-of-Science and Google Scholar. Specifically, we used the following search term in each database on 4/30/2017: *(“face” OR “embodied” OR “mirroring” OR “action perception” OR “action execution” OR “imitation” OR “empathy” OR “emotion recognition” OR “theory of mind” OR “mentalizing”) AND (“white matter” OR “tract” OR “pathway” OR “structural connectivity” OR “anatomical connectivity”) AND (“imaging” OR “MRI” OR “diffusion” OR “dMRI” OR “DTI” OR “tractography” OR “structural MRI” OR “morphometry” OR “direct electrical stimulation” OR “brain stimulation”)*. It is important to note that the initial search resulted in numerous clinical studies on social disorders that revealed abnormalities in WM structures in a patient group in comparison to a healthy cohort. However, making a simple comparison between patients with a social disorder and a healthy group is not enough to establish specific associations between WM and social cognition, because the observed WM differences could be caused by patients’ non-social deficits or symptoms (e.g. autism) (Travers et al., 2012). Therefore, we excluded studies with simple patient-control comparisons and only included papers with correlation analyses between WM and social cognition measures (see Figure 3 for paper selection procedure). This yielded a final sample of 51 studies on 3745 subjects (see Tables 1-3 for details).

**Figure 3.**
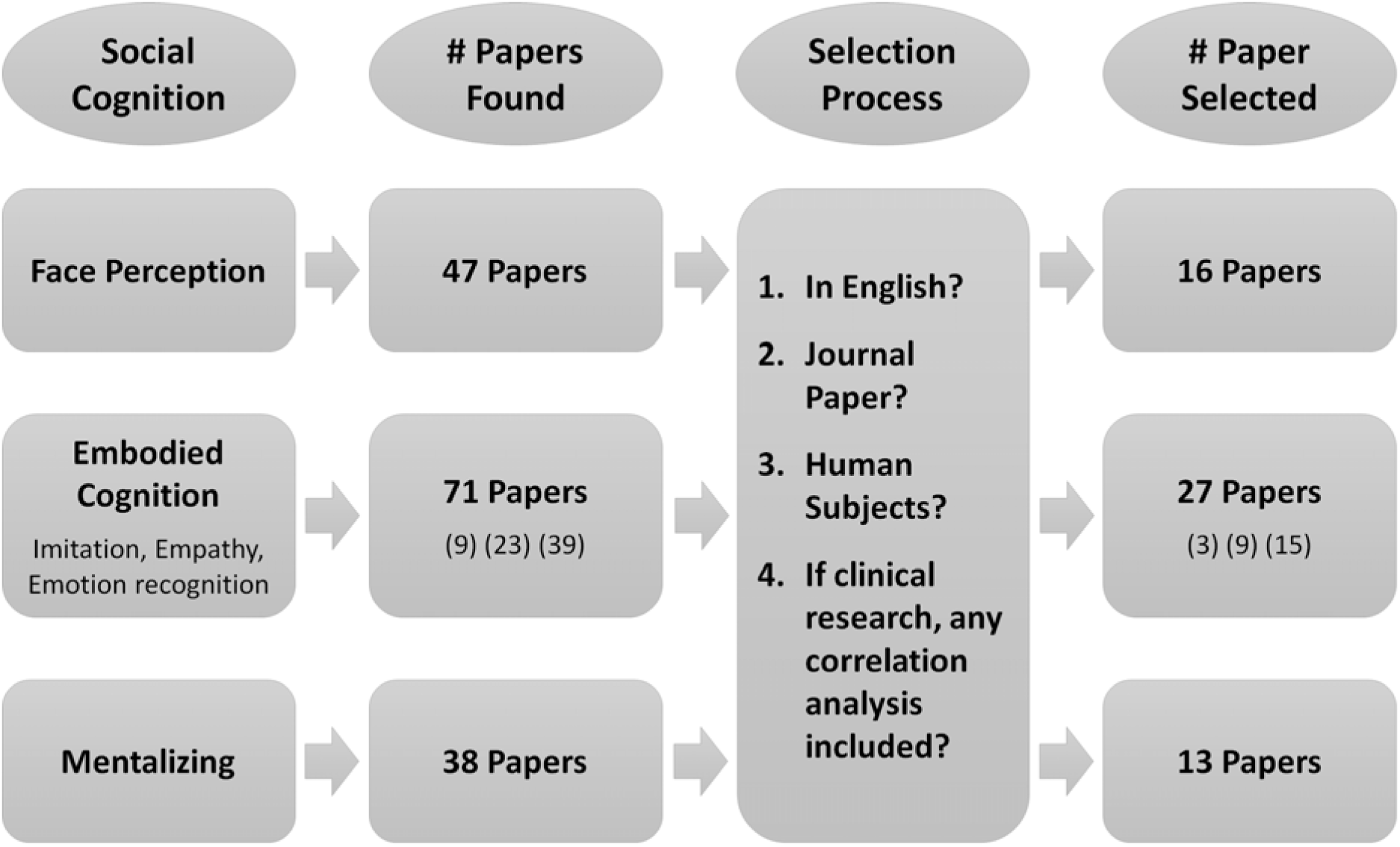
The literature review procedure, the inclusion criteria, and the number of surveyed studies for each network.

**Table 1.** Summary of 16 studies linking white matter to the face perception network.

| Study | Modality | Gradient Directions /B-value | Analysis Methods | Participants (N) | Mean Age | Social Cognition Measurements | Associated Tracts | Findings |
| --- | --- | --- | --- | --- | --- | --- | --- | --- |
| Ethofer (2011) | DTI fMRI | 30/1000 | Probabilistic Tractography | healthy adults (N=22) | 26 | gaze perception task (fMRI) | right SLF | The posterior STS and AI are anatomically connected by the SLF and their functional connectivity is modulated by gaze shifts. |
| Ethofer (2013) | DTI fMRI | 30/1000 | Probabilistic Tractography | healthy adults (N=23) | 23 | face localizer (fMRI) | right SLF | Face-selective STS anatomically connects with anterior face areas (IFG or OFC) via the dorsal SLF. |
| Gschwind (2012) | DTI fMRI | 30/1000 | Probabilistic Tractography | healthy adults (N=22) | 28 | face localizer (fMRI) | right ILF, right IFOF, right SLF | The study reveals the WM connectivity between face-responsive regions (OFA, FFA, AMY and STS). Strong WM connectivity is found for EVC-OFA, OFA-FFA and EVC-AMY, all via ILF (with right hemisphere predominance). AMY has stronger connections with EVC than OFA/FFA. STS is not connected to FFA/OFA, but rather lined to more frontal regions via AF/SLF |
| Marstaller (2016) | DTI fMRI | 64/3000 | Probabilistic Tractography | healthy adults (N=28) | 26 | face expression matching task | Bilateral ILF | Facial expression recognition skills can be predicted by the structural connectivity between FFA and AMG. People who are faster at discriminating threat-related expressions showed higher FA in this pathway (via ILF) |
| Iidaka (2012) | DTI fMRI | 64/3000 | Probabilistic Tractography | healthy adults (N=30) | 22 | face localizer (fMRI); Autism quotient (AQ) | right ILF | The volume of STS-AMG connectivity is positively correlated with the AQ score in the healthy sample. |
| Pyles (2013) | dMRI-NT fMRI | 257/7000 | Deterministic Tractography | healthy adults (N=5) | 28 | face localizer (fMRI) | right ILF, right IFOF | The study reveals the WM connectivity between face-responsive regions (OFA, FFA, ATL and STS). Strong WM connectivity were found for OFA-FFA, FFA-ATL and OFA-ATL, all via ILF and IFOF (with right hemisphere predominance). No connection was found for STS-OFA, STS-FFA and STS-ATL |
| Saygin (2012) | DTI fMRI | 60/700 30/700 | Probabilistic Tractography | healthy adults (N=23) | 28 | face localizer (fMRI) | N.A. | Anatomical connectivity patterns predict face selectivity in the FFA. |
| Scherf (2014) | DTI fMRI | 6/850 | Deterministic Tractography | healthy subjects aged 6-23 years (N=50) | 16 | face localizer (fMRI) | right ILF | The volume of FFA activation is positively correlated with the volume of right ILF, but negatively correlated with the MD in right ILF. |
| Tavor (2014) | DTI | 19/1000 | ROI-based, Deterministic Tractography | healthy adults (N=22) | 25 | face memory test | right ILF | Face memory accuracy is negatively correlated with FA (and positively correlated with RD) in the anterior portion of right ILF. |
| Unger (2016) | DTI | 64/1000 | Deterministic Tractography | healthy adults (N=28) | 22 | Cambridge task; Philadelphia face matching task; | right ILF, bilateral IFOF | Face memory accuracy is negative correlated with FA in right ILF and right IFOF, but positively correlated with FA in the left IFOF |
| Gomez (2015) | DTI<br>fMRI | 30/900<br>96/2000 | ROI-based, Probabilistic Tractography | developmental prosopagnosia patients (DP N=8; control N=18) | P=34<br>C=26 | face localizer (fMRI); Benton test; Cambridge task; | local right FFA fibers (ILF is not involved) | Healthy controls show a positive correlation between FA in local WM of right FFA and face recognition and memory performance, whereas DP patients show a negative correlation. DP patients also exhibit lower MD values in local WM of FFA in both hemispheres. |
| Grossi (2014) | DTI | 32/1000 | Deterministic Tractography | progressive prosopagnosia patients (PP N=1; control N=7) | P=65<br>C=68 | Benton test; Celebrity face recognition test; | right ILF | PP patients have a severe reduction of fibers (number of streamlines) in right ILF versus left ILF, while IFOFs are relatively symmetric. They also have higher MD in right ILF than controls. |
| Song (2015) | DTI<br>fMRI | 61/1000 | Voxel-based, Deterministic Tractography, Probabilistic Tractography | developmental prosopagnosia patients (DP N=16; control N=16) | P=31<br>C=30 | face localizer (fMRI); Cambridge task; celebrity face recognition test | local FFA fibers, bilaterally | DP patients show WM deficits only in local FFA (lower FA values), not in long-range tracts (IFOF and ILF). For both patients and controls, FA and MD in local FFA WM are correlated with face recognition performance. |
| Thomas (2008) | DTI | 6/850 | Deterministic Tractography | healthy adults across lifespan (N=28) | 42 | face matching task | right IFOF | A decrease in FA and volume in right IFOF by aging process is associated with a decrease in face discrimination accuracy |
| Thomas (2009) | DTI | 6/850 | Voxel-based, Deterministic Tractography | developmental prosopagnosia patients (DP N=6; control N=17) | P=58<br>C=56 | celebrity face recognition test | right ILF, right IFOF | DP patients have lower FA in bilateral ILF and IFOF, compared to controls. For both groups, the FA and number of streamline in right ILF and IFOF are positively correlated with face recognition performance. |
| Valdés-Sosa (2011) | DTI<br>sMRI<br>fMRI | 12/1200 | Deterministic Tractography, Probabilistic Tractography | acquired prosopagnosia patients (AP N=1; control N=10) | P=73<br>C=70 | neuropsychological tests on prosopagnosia | right ILF, right IFOF | This study reported an AP case who had disrupted FFA and ILF, but preserved anterior face-selective areas as well as covert face recognition skills. It suggests covert face skills might be subserved by the preserved IFOF connecting OFA to anterior face-selective areas. |

dmMRI-NT: diffusion-weighted MRI with non-tensor modeling; EVC: early visual cortices. Other acronyms can be referred to from the abbreviations section in the main text.

**Table 2.** Summary of 27 studies linking white matter to the mirroring network.

| Study | Modality | Gradient Directions /B-value | Analysis Methods | Participants (N) | Mean Age | Social Cognition Measurements | Associated Tracts | Findings |
| --- | --- | --- | --- | --- | --- | --- | --- | --- |
| **Imitation** |  |  |  |  |  |  |  |  |
| Hamzei (2016) | DTI fMRI | 61/1000 | Probabilistic Tractography | healthy adults (N=116) | 26 | imitation tasks (fMRI) | left SLF/AF, left extreme capsule | There are two anatomical pathways between frontal and parietal mirroring areas: the SLF III and AF (the dorsal stream) and the extreme capsule (the ventral stream) |
| Hecht (2013) | DTI | 60/1000 | Probabilistic Tractography | healthy adults (N=30) | 20 | N.A. | SLF, extreme capsule, ILF, all bilateral | In humans, SLF III connects IFG and IPL; SLF III, AF and extreme capsule connect IFG and STS; ILF connects IPL and STS. |
| Fishman (2015) | DTI fMRI | 61/1000 | Probabilistic Tractography | Children with autism (ASD N=35; control N=35) | P=14<br>C=13 | N.A. | left SLF | For ASD children, reduced FA and increased MD are found in WM tracts within the mirroring network (i.e. SLF) and this reduced WM integrity is correlated with individual ASD symptom severity. |
| **Emotion Recognition** |  |  |  |  |  |  |  |  |
| Ethofer (2012) | DTI fMRI | 30/1000 | Probabilistic Tractography | healthy adults (N=22) | 26 | passively listen to emotional voices (fMRI) | bilateral SLF and extreme capsule | Vocal emotion recognition is subserved by bilateral SLF and extreme capsule connecting IPL and IFG. |
| Ethofer (2013) | DTI fMRI | 30/1000 | Probabilistic Tractography | healthy adults (N=23) | 23 | emotional recognition task (fMRI) | right SLF | Face/voice emotion recognition engages posterior STS and IFG, and these two regions are connected by dorsal SLF. |
| Taddei (2012) | DTI EEG | 35/1000 | ROI-based, Deterministic Tractography, TBSS | healthy children (N=20) | 15 | passive facial emotion observation (EEG) | left UF, bilateral ILF and IFOF | N400 amplitudes in response to angry faces are negatively correlated with FA in left UF and bilateral ILF and IFOF. |
| Unger (2016) | DTI | 64/1000 | Deterministic Tractography | healthy adults (N=28) | 22 | facial emotion recognition task | right ILF, bilateral IFOF | The AD in right ILF and the FA in bilateral IFOF are negatively correlated with face emotion recognition performance. |
| Baggio (2012) | DTI | 30/1000 | ROI-based, TBSS | patients with Parkinson's disease (PD N=39; control N=23) | P=63<br>C=61 | Ekman facial emotion recognition | right IFOF, left IFOF/ILF, CC | For PD patients, FA in three WM tracts are positively correlated with the sadness identification performance: frontal portion of right IFOF (near IFG), left ILF/IFOF (near STS) and CC (right forceps minor). |
| Crespi (2014) | DTI | 32/1000 | ROI-based, TBSS | patients with amyotrophic lateral sclerosis (ALS N=22; control N=55) | P=60<br>C=62 | Ekman facial emotion recognition | right IFOF, right ILF | For ALS patients, the accuracy of facial emotion recognition (especially negative emotions) shows a positive correlation with FA values of both the right ILF and IFOF. |
| Crespi (2016) | DTI | 32/1000 | TBSS | patients with amyotrophic lateral sclerosis (ALS N=13; control N=14) | P=59<br>C=56 | story-based emotion attribution task | right IFOF and UF, left SLF and CC | For ALS patients, emotion attribution scores are positively correlated with FA in right IFOF and UF, left SLF, CC (genu and forceps minor). |
| Downey (2015) | DTI | 64/1000 | TBSS | patients with behavioral variant frontotemporal dementia (bvFTD N=44; control N=37) | P=64<br>C=63 | the awareness of social inference test | right ATR and UF, fornix | For bvFTD patients, emotion identification score is negatively correlated with AD, RD, and MD in fornix and right ATR; sarcasm identification score is negatively correlated with AD, RD, and MD in right UF. |
| Fujie (2008) | DTI | 12/700 | Deterministic Tractography | patients with mild cognitive impairment (MCI N=16; control N=16) | P=71<br>C=71 | facial emotion recognition task | left UF | For MCI patients, FA values of the left UF are positively correlated with the performance of face emotional recognition for sadness, fear, and surprise. |
| Genova (2015) | DTI | 12/1000 | TBSS | patients with traumatic brain injury (TBI N=42; control N=23) | P=35<br>C=39 | facial emotion identification test | right ILF and IFOF | For TBI patients, the face emotion discrimination score is positively correlated with FA in right IFOF, but negatively correlated with AD, MD, and RD in right ILF. |
| Philippi (2009) | sMRI | N.A. | Voxel-based , ROI-based | patients with brain-damage (N=103; control N=18) | P=52<br>C=56 | facial emotion recognition task | right ILF, IFOF and SLF | Damage to the right IFOF, ILF and SLF predicts facial emotion recognition impairment. Damage to right IFOF specifically impairs fear recognition |
| Jalbrzikowski (2014) | DTI | 64/1000 | ROI-based, TBSS | patients with velocardiiofacial syndrome (VCFS N=36; control N=29) | P=16<br>C=16 | Penn emotion recognition test | left IFOF and UF. | For VCFS patients, AD in the left IFOF and UF is positively correlated with emotional recognition performance, especially fear recognition. |
| Radoeva (2012) | DTI | 15/800 | ROI-based | patients with velocardiiofacial syndrome (VCFS N=33; control N=16) | P=18<br>C=18 | facial emotion recognition task | right SLF and IFOF | For VCFS patients, emotion recognition score is positively correlated with AD in right SLF and right IFOF. |
| Mike (2013) | sMRI | N.A. | Voxel-based | patients with multiple sclerosis (MS N=49; control N=24) | P=40<br>C=37 | facial emotion recognition test | left fornix; right ILF and IFOF, bilateral UF; CC | For MS patients, facial emotion recognition performance is negatively correlated with lesion volume in genu and splenium of CC, right ILF and IFOF, bilateral UF, and left fornix. |
| Saito (2017) | DTI | 51/900 | Deterministic Tractography | patients with schizophrenia (SZ N=16; control N=16) | P=20<br>C=22 | perception of emotional intimacy | left SLF | For SZ patients, MD in IPL-IFG connection (via SLF) is positively correlated with the disrupted perception of emotional intimacy (e.g. inability to feel intimacy). |

| Empathy |  |  |  |  |  |  |  |  |
| --- | --- | --- | --- | --- | --- | --- | --- | --- |
| Chou (2011) | DTI | 13/900 | TBSS | healthy adults (N=80) | 26 | empathy quotient (EQ) | local WM in left IPL and STS; right ATR and SLF; left ILF | WM underlying empathy is sex-dependent. EQ score is positively correlated with FA in multiple regions (left IPL and STS) and tracts (right ATR and SLF, left ILF) in females, but negatively correlated with FA in these regions/tracts in males. |
| Mueller (2013) | DTI<br>sMRI<br>fMRI | 20/1000 | TBSS | patients with high-functioning autism (ASD N=12; control N=12) | P=36<br>C=33 | questionnaires of empathy and emotionality | N.A. | WM in local TPJ is correlated with emotionality. But no measures correlate with empathy scores |
| Nakagawa (2015) | DTI<br>sMRI | 32/1000 | Voxel-based<br>ROI-based | healthy adults (N=776) | 20 | empathy quotient (EQ) | local WM in bilateral IPL, right AI/IFG, and left TPJ/STS | For lonely individuals, the EQ score is positively correlated with local WM density in bilateral IPL, right AI/IFG, and left TPJ/STS |
| Parkinson (2014) | DTI | 32/1000 | TBSS | healthy adults (N=64) | 19 | interpersonal reactivity index | IFOF, SLF, UF, ATR, CC, CST, all bilaterally; right ILF | The 'empathic concern' subscales of emotional empathy is positively correlated with FA in right ILF, bilateral SLF, IFOF, UF, ATR, CST, and CC (forceps minor) |
| Takeuchi (2013) | DTI<br>sMRI | 32/1000 | Voxel-based | healthy adults (N=567) | 21 | empathy quotient (EQ) | local WM in social regions, left SLF, ILF and IFOF, fornix, CC | EQ score is positively correlated with local WM volume in multiple regions (right IFG, IPL, TPJ, PCC, and MPFC) and tracts (left ILF, left IFOF, fornix, genu of CC). The EQ score is also positively correlated with FA in left SLF. |
| Fujino (2014) | DTI | 81/1500 | TBSS | patients with schizophrenia (SZ N=69; control N=69) | P=37<br>C=34 | interpersonal reactivity index (IRI) | CC, left IFOF, left ATR | For SZ patients, the 'personal distress' subscales of empathy are negatively correlated with FA in the splenium of the CC, and the 'fantasy' subscales are positively correlated with FA in the left IFOF and ATR. |
| Herbet (2015) *Neuro psychologia* | sMRI | N.A. | Voxel-based, ROI-based | patients with surgical resection for diffuse low-grade glioma (DLGG N=107) | 41 | empathy quotient (EQ) | right UF and IFOF | For DLGG patients, disconnection of right UF predicts a low subjective empathy and disconnection of right IFOF predicts a high subjective empathy. |
| Oishi (2015) | DTI<br>sMRI | 6/1000 | ROI-based | patients with acute ischemic stroke (N=30) | N.A. | emotional empathy task | right UF | Damage to the right UF is negatively correlated with emotional empathy performance. |
| Olszewski (2017) | dMRI-NT | 64/700 | Deterministic Tractography | patients with velocardiofacial syndrome (VCFS N=57; control N=30) | P=21<br>C=21 | trait emotional intelligence questionnaire (empathy subsets) | right ATR and UF | For VCFS patients, empathy scores is positively correlated with RD in right ATR and negatively correlated with the number of streamlines in right UF. |

bvFTD: behavioral-variant frontotemporal dementia; DLGG: diffuse low-grade glioma; dMRI-NT: diffusion-weighted MRI with non-tensor modeling; TBI: traumatic brain injury; SZ: schizophrenia; VCFS: velocardiofacial syndrome. Other acronyms can be referred to from the abbreviations section in the main text.

**Table 3.**
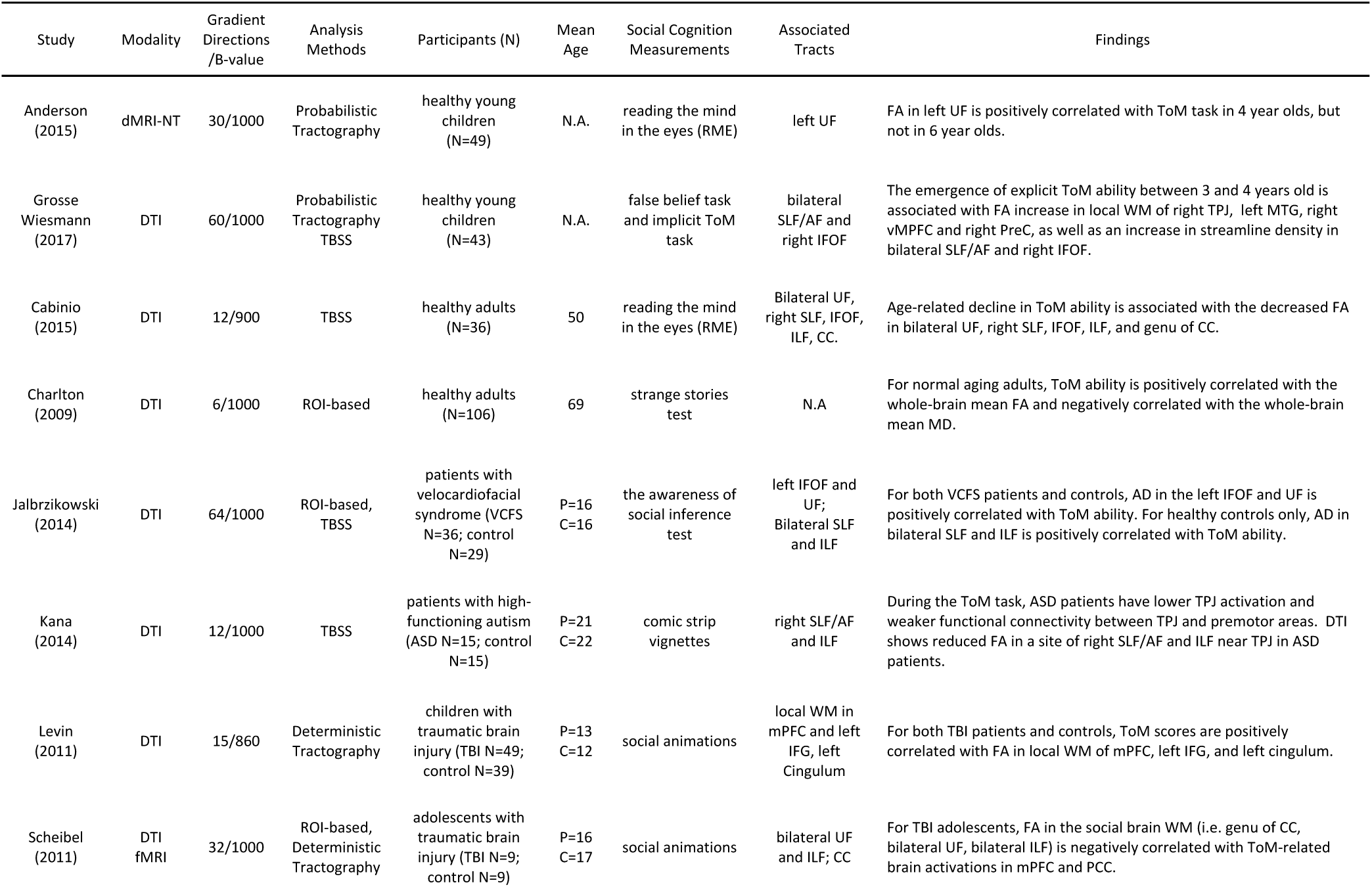
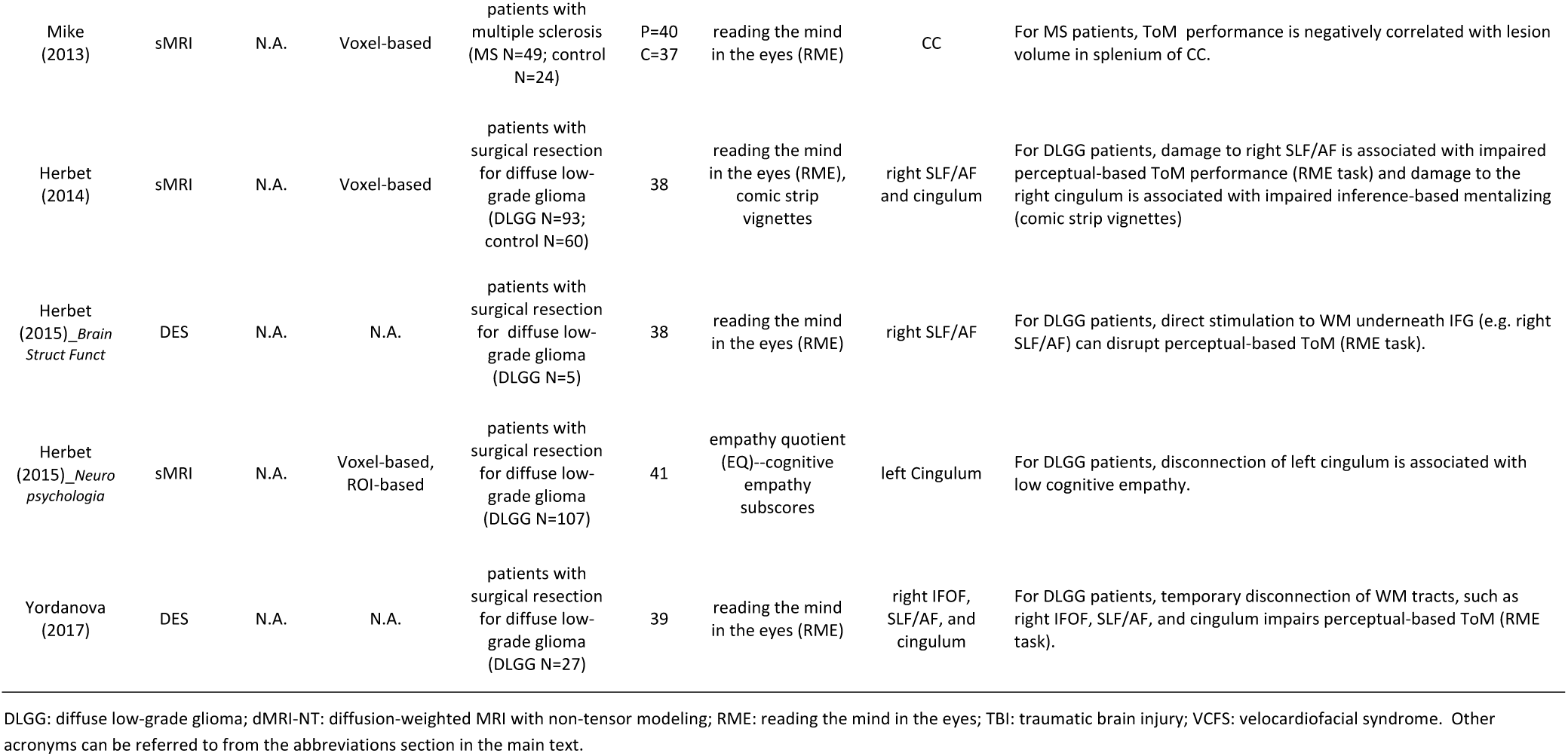
Summary of 13 studies linking white matter to the mentalizing network.

### Methodological Summary

Figure 4 summarizes a couple of key features in the literature. As can be seen, the sample size varies across studies (ranging from 5 to 766) and depends on technique modality. While most sMRI research tended to recruit large numbers of subjects (>80), dMRI studies typically had a moderate sample size (20 to 40). Although DES studies had the smallest sample size, their findings should not be discounted, because the technique is more rigorous and reliable than dMRI and sMRI. In addition, it is clear that the most common data analysis method was tractography-based and the most common dMRI measure was FA, although many studies used overlapping measures and methods. Figure 4 also illustrates the frequency of two critical data acquisition parameters used in dMRI research: the “gradient directions” and the “b-value”. The number of diffusion-encoding gradient directions defines the number of orientations at which diffusion signals are sampled. As the number increases, more diffusion-weighted images are used for the calculation of the diffusion tensor model, resulting in more accurate estimation of microstructural indices related to the tensor; however, a large number of gradient directions substantially elongates the required scanning time. The b-value represents the degree of diffusion weighting and determines the strength and duration of the diffusion gradients. The ability to delineate WM fasciculi oriented in different directions improves as the b-value increases, but higher b-values (e.g. >3000) come at a cost of lower signal-to-noise ratio (Jones et al., 2013). As we can see from Figure 4, most dMRI studies in the literature used 17-32 gradient directions with a b-value of 1000 s/mm^2^.

**Figure 4.**
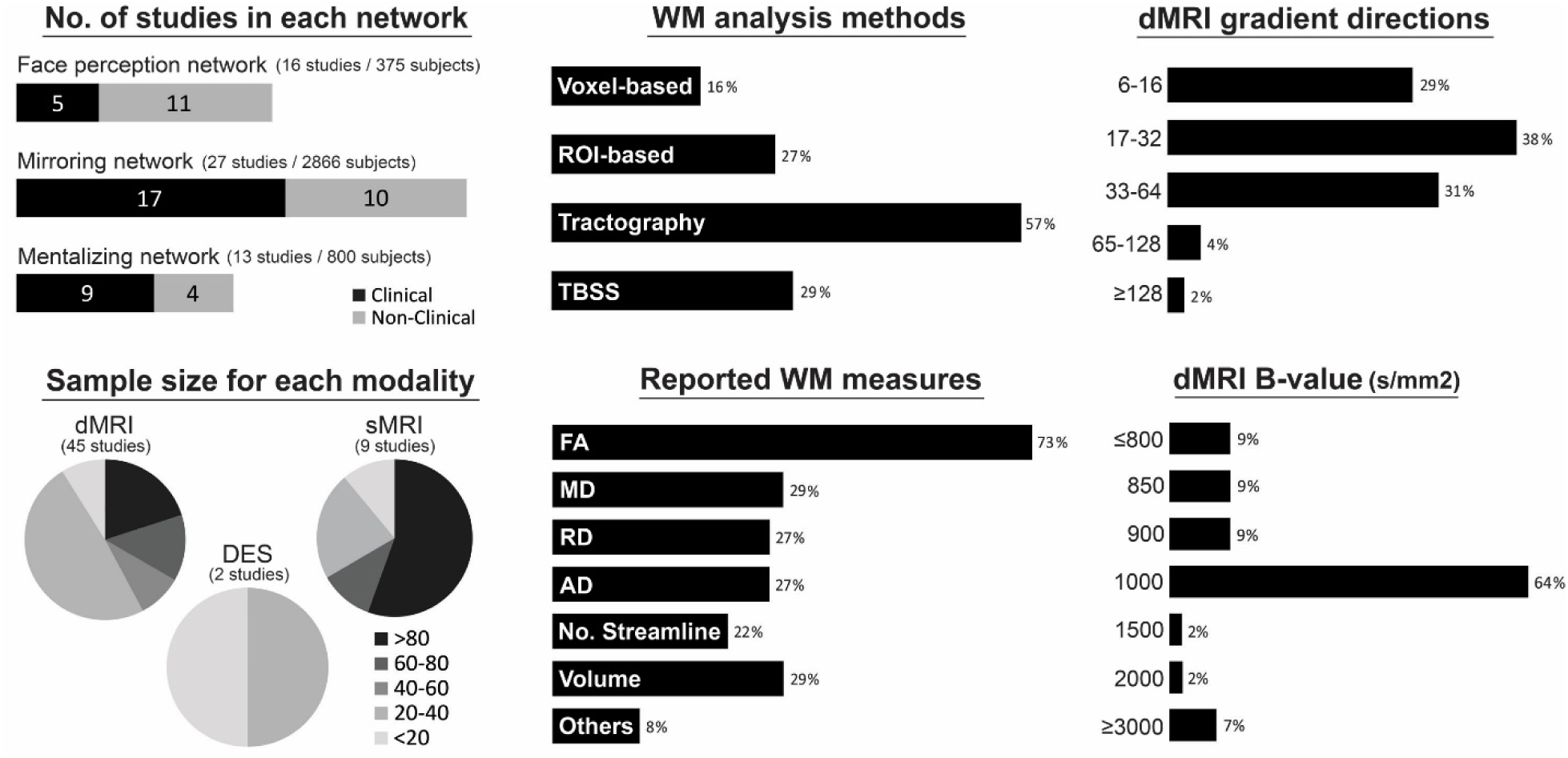
Key features of the 51 empirical studies surveyed in the present paper, including number of clinical/non-clinical studies for each social brain network, the sample size for each technique modality, how WM measures were analyzed and reported in the studies, and the diffusion data acquisition parameters (the gradient directions and b-values). Note: Percentage might add up to more than 100% because of studies often using more than one type of analysis method, measure, or acquisition protocol. AD, axial diffusivity; DES, direct electrical stimulation; dMRI, diffusion magnetic resonance imaging; FA, fractional anisotropy; MD, mean diffusivity; No. Streamline, number of streamline; RD, radial diffusivity; sMRI, structural magnetic resonance; TBSS, tract-based spatial statistics;

We found that more than half of the studies, particularly of the mirroring and mentalizing network, are based on clinical populations. They include major psychiatric and neurological disorders characterized by prominent social impairments, such as autism spectrum disorder (ASD) (Fishman, Datko, Cabrera, Carper, & Müller, 2015; Kana, Libero, Hu, Deshpande, & Colburn, 2014; Mueller et al., 2013), behavioral-variant frontotemporal dementia (Downey et al., 2015), and prosopagnosia (Gomez et al., 2015; Grossi et al., 2014; Song et al., 2015; Thomas et al., 2008, 2009; Valdés-Sosa et al., 2011), as well as those with secondary impairments in social cognition, such as schizophrenia (Fujino et al., 2014; Saito et al., 2017), amyotrophic lateral sclerosis (ALS) (Crespi et al., 2014, 2016), mild cognitive impairment (MCI) (Fujie et al., 2008), traumatic brain injury (Genova et al., 2015; Levin et al., 2011; Scheibel et al., 2011), Parkinson’s disease (PD) (Baggio et al., 2012), brain lesion/stroke (Oishi et al., 2015; Philippi et al., 2009), velocardiofacial syndrome (Jalbrzikowski et al., 2014; Olszewski et al., 2017; Radoeva et al., 2012), multiple sclerosis (MS) (Mike et al., 2013) and diffuse low-grade glioma (Herbet et al., 2014; Herbet, Lafargue, Moritz-Gasser, Menjot de Champfleur, et al., 2015; Herbet, Lafargue, Moritz-Gasser, Bonnetblanc, & Duffau, 2015; Yordanova, Duffau, & Herbet, 2017). In terms of social cognitive measurements, the literature has employed several behavioral paradigms to probe each social function (see Table 1-3). For example, face processing skills were measured by celebrity face recognition tasks (Grossi et al., 2014; Song et al., 2015; Thomas et al., 2009), face matching tasks (e.g. Benton tests, Philadelphia battery) (Gomez et al., 2015; Grossi et al., 2014; Thomas et al., 2008; Unger et al., 2016), and face memory tasks (e.g. Cambridge tests) (Gomez et al., 2015; Song et al., 2015; Unger et al., 2016); empathy was assessed by the “empathy quotient” (Chou et al., 2011; Herbet, Lafargue, Moritz-Gasser, Menjot de Champfleur, et al., 2015; Nakagawa et al., 2015; Takeuchi et al., 2013), the “interpersonal reactivity index” (Fujino et al., 2014; Parkinson & Wheatley, 2014), and the “trait emotional intelligence questionnaire” (empathy subsets) (Olszewski et al., 2017); mentalizing abilities were evaluated by “false belief” stories (Grosse Wiesmann et al., 2017), cartoon animations (Levin et al., 2011; Scheibel et al., 2011), comic strip vignettes (Herbet et al., 2014; Kana et al., 2014), and the “reading the mind in the eyes” task (Anderson et al., 2015; Cabinio et al., 2015; Herbet et al., 2014; Herbet, Lafargue, Moritz-Gasser, Bonnetblanc, et al., 2015; Mike et al., 2013; Yordanova et al., 2017). Such a wide variety of seemingly disparate disorders as well as diverse behavioral paradigms provide excellent opportunities for exploring the relationship between WM tracts and social functions.

### Major Findings

#### Face Perception Network

Two WM tracts are repeatedly reported in the face perception literature: the inferior longitudinal fasciculus (ILF) and the inferior fronto-occipital fasciculus (IFOF) (see Table 1). They are the main associative bundles that project through occipito-temporal cortex, connecting the occipital lobe to the temporal, and frontal lobes, respectively (Rokem et al., 2017). The ILF is a monosynaptic pathway connecting ventral extrastriate regions, and in some cases portions of the inferior parietal lobe, to the anterior temporal lobe, the hippocampus, and the amygdala (Catani, Jones, Donato, & Ffytche, 2003). The IFOF begins in the ventral occipital cortex, continues medially through the temporal cortex dorsal to the uncinate fasciculus, and terminates in the inferior frontal, medial prefrontal, and orbitofrontal cortex (Catani & Thiebaut de Schotten, 2008). dMRI tractography combined with functional face localizer confirmed that the ILF connects multiple pairs of face perception network nodes including the OFA-FFA, OFA-ATL, FFA-ATL, FFA-AMG and STS-AMG (Gschwind et al., 2012; Iidaka et al., 2012; Pyles et al., 2013), while the IFOF connects the OFA-IFG (Valdés-Sosa et al., 2011). Converging evidence, described below, indicates that these two tracts are critically important for face processing.

First, the early development of the right ILF is associated with the emergent functional properties of the face perception network. Using both DTI and fMRI, Scherf et al., (2014) investigated whether developmental differences in the structural properties of bilateral ILF were related to developmental differences in the functional characteristics of the face-processing regions connected by ILF (e.g. OFA, FFA). Across children, adolescents, and adults (ages 6–23 years), they found bilateral ILF exhibited an age-related increase in volume, and those individuals with larger right ILF volumes also exhibited larger right FFA volumes. This suggests a tight relationship between the structural refinements of right ILF and functional selectivity in the developing face perception network.

Similarly, age-related declines in face perception skills have been linked to degeneration of the right IFOF. Thomas and colleagues (2008) used DTI to scan subjects across a wide age range (18-86 years) and also measured individual performance on face perception tasks. They observed that the right IFOF was the only tract that decreased in volume as a function of age, and subjects with smaller volumes and lower FA values in the right IFOF exhibited worse performance on the face matching task. This evidence indicates that the right IFOF is vulnerable to the aging process, and age-related decreases in the structural properties of this tract might be responsible for decrements in face processing abilities in aging adults.

Moreover, disruptions in ILF and IFOF are associated with face blindness in prosopagnosia. Developmental prosopagnosia (DP) is a social disorder characterized by a lifelong impairment in face recognition despite normal sensory vision and intelligence. Interestingly, DP patients exhibit normal patterns of fMRI activation in response to faces in posterior parts of the face perception network (e.g. OFA, FFA; Behrmann et al., 2005), but reduced activation in anterior nodes (e.g. ATL; Avidan et al., 2014). Based on this observation, it had been suggested that the impairments in DP might arise, not from a dysfunction of cortical parts of the face perception network, but from a failure to propagate signals from the intact posterior components to the compromised anterior components of the network. As the two major tracts that project through the posterior to anterior regions of the face network, the ILF and IFOF are top candidates for testing. As such, Thomas et al., (2009) scanned a group of DP patients and measured the severity of their face recognition deficits. Relative to the control group, the integrity in the right ILF and IFOF in DP patients was remarkably compromised (i.e. lower FA and volume) and the extent of this compromise was correlated with individual face perception deficits. This finding was interpreted as evidence for DP as a “disconnection syndrome”, i.e. face blindness occurs because intact posterior face processing regions are unable to communicate via the ILF and IFOF with more anterior regions. Similar findings were also observed in other types of prosopagnosia where patients exhibited severe fiber reductions in the right ILF (Grossi et al., 2014; Valdés-Sosa et al., 2011).

Other face processing abilities are also related to the ILF and IFOF, although the underlying mechanisms are unclear. Unger et al., (2016) showed that face memory accuracy was negatively correlated with FA in the right ILF and IFOF, but positively correlated with FA in the left IFOF. Tavor et al., (2014) reported similar findings and further indicated that the anterior part of right ILF explained the most inter-individual variation of face memory performance. Moreover, individual differences in processing facial communicative signals can be predicted by the structural connectivity between face-selective areas (e.g. FFA or STS) and amygdala (AMG) via the ILF. People who are better at discerning threat-related facial expressions showed higher FA in the FFA-AMG connectivity (Marstaller, Burianova, & Reutens, 2016), and people who have better social communication skills had larger volumes of the STS-AMG pathway (Iidaka et al., 2012).

Aside from the ILF and IFOF, the superior longitudinal fasciculus (SLF) seems to be a third WM tract that subserves face processing. Anatomically, the SLF connects superior-posterior face-selective regions, such as the STS, with anterior-inferior face-selective regions (IFG and OFC) (Ethofer et al., 2013; Gschwind et al., 2012). Functionally, the SLF has been associated with gaze processing (Ethofer et al., 2011) and face-voice integration (Ethofer et al., 2013).

It is important to note that almost every reported WM correlate of face processing skills is in the right hemisphere. This lateralization of WM function is consistent with the significant right-hemisphere predominance in the face perception literature: fMRI studies typically show larger face activations in the right, relative to the left hemisphere, and behavioral studies show better performance for faces presented in the left than the right visual fields (Tavor et al., 2014). Some studies speculate that the left ILF is more specialized for face tasks requiring access to language, such as face naming, while the right ILF may have functions more aligned with strictly visuospatial functions, such as face discrimination (Unger et al., 2016).

#### Mirroring Network

Converging evidence suggests that the superior longitudinal fasciculus (SLF) is the most important WM tract for embodied social cognition (see Table 2). The SLF is a large association bundle composed of medial and lateral fibers connecting the frontal, parietal, and temporal lobes (Kamali, Flanders, Brody, Hunter, & Hasan, 2014). This WM tract has a known role in language and spatial attention (Merchant, 2011) and has recently been identified to be the main fiber pathway for the fronto-parietal mirroring network (Hamzei et al., 2016; Hecht et al., 2013; Iacoboni & Dapretto, 2006; Parlatini et al., 2017). Several studies indicate that the SLF is functionally associated with imitation, empathy, and emotion recognition abilities. For example, Hecht et al., (2013) found that the evolved imitation skills across species (macaques, chimpanzees, and humans) can be explained by increased SLF connections supporting the fronto-parietal mirroring network. The empathy quotient is positively correlated with FA values in the SLF bilaterally, most extensively in the right SLF (Chou et al., 2011; Parkinson & Wheatley, 2014; Takeuchi et al., 2013). The SLF is also associated with individual’s emotion recognition ability, regardless of whether the task is face-based (Crespi et al., 2014; Philippi et al., 2009; Radoeva et al., 2012), story-based (Crespi et al., 2016), or voice-based (Ethofer et al., 2012, 2013). When the right SLF is disrupted by brain lesion (Philippi et al., 2009) or psychiatric disorders (Crespi et al., 2014, 2016; Radoeva et al., 2012; Saito et al., 2017), the integrity of SLF is also positively correlated with patients’ emotion recognition skills.

Other robust associations between WM and embodied cognition have been identified in three limbic tracts: the uncinate fasciculus (UF), the anterior thalamic radiation (ATR), and the fornix. The UF is a hook-shaped ventral associative bundle that links medial temporal areas (e.g. ATL, AMG) to portions of frontal cortices (both medial and lateral OFC) (Catani & Thiebaut de Schotten, 2008). It has been linked to episodic memory, semantic memory, and social-emotional processing (Von Der Heide, Skipper, Klobusicky, & Olson, 2013). The ATR is a major projection from the thalamus, which carries reciprocal connections from the hypothalamus and limbic structures (e.g. AMG, hippocampus) to the prefrontal cortex and ACC (Catani, Dell’Acqua, & Thiebaut de Schotten, 2013). It has been primarily implicated in affective processing and emotion regulation (Downey et al., 2015). The fornix is a core limbic tract directly connecting the hippocampus to the mammillary bodies and hypothalamus. It is mainly involved in episodic memory and evaluative processing (Catani & Thiebaut de Schotten, 2008). Disruption of these limbic tracts has been commonly observed in clinical disorders, such as the behavioral variant frontotemporal dementia, mild cognitive impairment (MCI), and velocardiofacial syndrome (Daianu et al., 2016; Liu et al., 2017; Perlstein et al., 2014), and these patients typically exhibit severe impairments in empathy and emotion recognition abilities (Jalbrzikowski et al., 2012; Kessels et al., 2007; Lough et al., 2006; Spoletini et al., 2008). Abnormal diffusivity (AD, RD, and MD) has been reported in the right ATR, UF, and fornix in frontotemporal dementia patients and correlates with disrupted understanding of emotion and sarcasm (Downey et al., 2015). Reduced FA in the left UF in MCI patients correlates with impaired emotion recognition and expression (Fujie et al., 2008). For patients with velocardiofacial syndrome, one study reported that empathy scores correlated with RD in the right ATR and negatively correlated with the number of streamlines in right UF (Olszewski et al., 2017); the patients’ emotional recognition performance for fear expression was also positively correlated with AD in the left UF (Jalbrzikowski et al., 2014). In addition, for patients with multiple sclerosis, facial emotion recognition performance is negatively correlated with lesion volume in bilateral UF, as well as the left fornix (Mike et al., 2013). For patients with acute ischemic stroke or surgical resection for a diffuse low-grade glioma, disconnection of the right UF predicted low empathy ability (Herbet, Lafargue, Moritz-Gasser, Menjot de Champfleur, et al., 2015; Oishi et al., 2015). For schizophrenia patients, subscales of empathy positively correlated with FA in the left ATR (Fujino et al., 2014). Finally, the integrity of these three limbic tracts not only predicts socio-emotional functioning in pathological circumstances, but also in normal individuals. For instance, higher FA in the UF or ATR has been associated with higher levels of empathy among healthy individuals (Parkinson & Wheatley, 2014), and larger WM volume in the fornix is associated with higher empathy quotient scores (Takeuchi et al., 2013).

The ILF and IFOF also appear to be important for emotional recognition and empathy. Two prior studies with large samples of patients with focal brain lesions reported that damage to the right ILF or IFOF correlates with impairments in facial emotion recognition, more specifically in the recognition of fear, anger, and sadness (Genova et al., 2015; Philippi et al., 2009). For patients with Parkinson’s disease, decreased FA in bilateral IFOF and the left ILF was associated with impaired sadness identification performance (Baggio et al., 2012). Additionally, the inter-individual variation of emotion recognition and empathy abilities among healthy adults can be predicted by the microstructure of the right ILF and bilateral IFOF (Parkinson & Wheatley, 2014; Unger et al., 2016). Little is known, however, about how the IFOF and ILF are implicated in embodied cognition, because the core mirroring network (STS, IPL, and IFG) is usually thought to be part of the dorsal stream (Hamzei et al., 2016). Since the two tracts directly project from early visual cortices to affective mirroring areas (i.e. AMG, AI, and ACC for the IFOF; AMG for the ILF) (Gschwind et al., 2012; Sarubbo, De Benedictis, Maldonado, Basso, & Duffau, 2013), it is possible that they engage in rapid evaluative processing paralleled with the basic embodied simulation process to facilitate accurate recognition of emotions. Another possibility stems from the nature of the biased behavioral paradigm used in the literature: almost all emotion recognition tasks are face-based, such as the Ekman Face Test, and it is already established that the IFOF and ILF are essential for face processing (Rokem et al., 2017).

Interestingly, sex differences in empathy may be reflected in sex differences in WM microstructure of the aforementioned tracts. Research on the “empathizing-systemizing” theory (Baron-Cohen, 2009) suggests that females generally perform better on emotion recognition and empathy tasks, whereas males excel in mental rotation, spatial navigation, and mathematics (i.e. systemizing). Two studies have shown that empathizing skill is positively correlated with microstructure in bilateral SLF, right ATR, right fornix, left ILF, and left IFOF in females, but negatively correlated with microstructure in these tracts in males (Chou et al., 2011; Takeuchi et al., 2013).

#### Mentalizing Network

The literature suggests that the cingulum and a portion of the SLF, the arcuate fasciculus, are two pivotal WM tracts for mentalizing abilities (see Table 3). The cingulum is a large association fiber pathway that encircles the corpus callosum, going from the medial prefrontal cortex/anterior cingulate cortex through the posterior cingulate cortex/precuneus, and from there to the medial temporal structures proximal to the hippocampus. It is part of the limbic system and is broadly involved in attention, memory, and emotional processing (Catani & Thiebaut de Schotten, 2008). Given that the cingulum provides strong structural connections between the MPFC and PCC, it has been argued as the main structural skeleton of the default mode network (van den Heuvel, Mandl, Luigjes, & Hulshoff Pol, 2008) and the mentalizing network (Yordanova et al., 2017). The arcuate fasciculus (AF) has long been implicated in language processing, as it connects Wernicke’s area to Broca’s area in the left hemisphere; however, the function of the right AF remains unclear. It has recently been proposed that the right AF might subserve mentalizing (Herbet et al., 2014), since the tract connects frontal cortices with the right TPJ, a region responsible for thinking about others’ thoughts and intentions (Saxe & Wexler, 2005).

Several clinical studies have reported that mentalizing abilities are compromised when the cingulum or right AF is disrupted. For children with traumatic brain injury, the severity of the ToM impairment is positively correlated with the degree of axonal injury in the left cingulum (Levin et al., 2011). Individuals with high-functioning autism (ASD) had lower right TPJ activation, weaker functional connectivity between the TPJ and frontal areas during the ToM task, and most critically, reduced WM integrity in the right AF near the TPJ (Kana et al., 2014). Perhaps the most compelling evidence comes from two studies using direct electrical stimulation (DES) of WM tracts during neurosurgery—the only technique that allows for direct information on the functional role of WM tracts in cognition. Both studies found that virtual disconnection of the right AF or cingulum severely impairs the accuracy of mental state attribution (Herbet, Lafargue, Moritz-Gasser, Bonnetblanc, et al., 2015; Yordanova et al., 2017). This suggests that proper functioning of these tracts is essential for normal mentalizing abilities.

There is some evidence that these two tracts might be specialized for different mentalizing processes. Studies on patients with diffuse low-grade gliomas revealed that damage to the right cingulum is associated with impaired performance on inference-based tasks (e.g. comic strip vignettes), whereas damage to the right AF is associated with impaired performance on perceptual-based ToM tasks, such as tested in “reading the mind in the eyes” (Herbet et al., 2014). Considering that the cingulum and AF connect different nodes of the mentalizing network (the cingulum mainly projects to medial nodes, such as MPFC and PCC, while the AF projects to lateral nodes, such as the TPJ and IFG), this double dissociation in terms of WM function resonates with previous fMRI studies showing that the MPFC engages most in inference-based ToM tasks, whereas the IFG only activates during perceptual-based ToM tasks (Schurz et al., 2014).

Substantial evidence in fMRI research suggests a critical role of the amygdala in ToM, especially for face-based mental state inferences (Mar, 2011). This may be due to the amygdala’s role in guiding attention to the eye region of the face, which may be an important first step in the process of interpreting the mental states of others (Adolphs & Spezio, 2006). However, the amygdala does not operate in isolation: WM tracts connecting the amygdala to other mentalizing areas may also contribute to ToM processes. Several studies have shown that amygdala-related WM tracts (i.e. UF, IFOF, and ILF) are important for accurate mentalizing. For example, impaired ToM skills in patients with velocardiofacial syndrome are associated with WM microstructural alterations in the left IFOF, left UF, and bilateral ILF (Jalbrzikowski et al., 2014). For patients with surgical resection due to the presence of gliomas, transient disconnection of the right IFOF by DES impairs performance on the “reading the mind in the eyes” task. Cross-sectional research also supports the crucial role of these amygdala-related WM tracts in lifespan changes in ToM abilities. Using TBSS, Grosse Wiesmann et al., (2017) found that the emergence of explicit ToM abilities between 3 and 4 years of age is associated with an increase in streamline density in the right IFOF and bilateral SLF/AF. Another study revealed that variation in the microstructure of the left UF positively correlates with inter-individual variance of “reading the mind in the eyes” task performance in 4-year-olds, but not in 6-year-olds, suggesting that the UF might be more important for the emergence, but not maintenance, of ToM function (Anderson et al., 2015). In addition, age-related declines in ToM abilities throughout the lifespan have been associated with decreased FA in bilateral UF, right IFOF, and right SLF (Cabinio et al., 2015).

### Summary

To summarize, three major tracts in the right hemisphere have been implicated in face processing: the ILF, the IFOF, and the SLF. Studies in young children and older adults, as well as patients with prosopagnosia, all attest to the crucial role these tracts play in skilled face perception. The literature on imitation, empathy, and emotional recognition identifies the SLF as the most critical tract for embodied social processes, and it has also been identified as the primary fiber pathway for the mirroring network. The UF, ATR, and fornix have also shown robust associations with embodied social processes. Disruption of these limbic tracts causes severe impairments in empathy and emotion recognition abilities across a variety of clinical disorders. Finally, WM research on ToM suggests that the cingulum and the AF are essential for mentalizing abilities. This claim is bolstered by strong evidence from DES studies. Additionally, changes in ToM abilities across the lifespan are associated with amygdala-related WM tracts (i.e. UF, IFOF, and ILF). Bear in mind that our literature review tries to draw conclusions more generally from the entire body of WM studies, rather than from any single finding.

It is also worth noting that healthy WM in the corpus callosum (CC) appears to be important for social cognition, as our literature review shows apparent involvement of the CC in both embodied cognition (Baggio et al., 2012; Crespi et al., 2016; Fujino et al., 2014; Mike et al., 2013; Parkinson & Wheatley, 2014; Takeuchi et al., 2013) and ToM (Cabinio et al., 2015; Mike et al., 2013; Scheibel et al., 2011). This is consistent with research on autism and agenesis of the corpus callosum, which both reveal that corpus callosum abnormalities can cause severe impairments in social functioning in the real world (Paul et al., 2007; Travers et al., 2012). One appealing hypothesis (Kennedy & Adolphs, 2012) is that social cognition is contingent upon rapid and reliable communication between social brain areas that are spatially separate, such as language-related areas in the left hemisphere and face processing areas in the right hemisphere. Given the highly interactive, real-time nature of social behavior, there is substantial pressure to integrate contralateral processing as efficiently as possible; therefore, social cognition requires considerable amounts of myelinated corpus callosum connections across hemispheres.

## Elucidating Anatomical Architecture of Social Brain Networks

The above literature review has informed us of several important WM tracts for social cognition. However, the exact architecture of interconnections between social brain regions still remains unknown. Unraveling this connectivity profile is extremely useful when we speculate or interpret results, because once we find a correlation between a WM tract (e.g. UF) and a social behavior/disorder, we would like to infer what underlying neural communications (e.g. AMG-MPFC interaction) are potentially involved or disrupted. A second motivation is to bridge the conceptual gap between two major analysis methods used in the DTI literature. The TBSS method tends to report findings based on the tract name listed in a standard brain atlas (e.g. SLF, ATR), whereas the tractography-based studies frequently report results in terms of pathways and seed ROIs (e.g. STS-IFG pathway). It is difficult to compare findings from these two methods without knowing the tract composition of each pathway. Last, sample sizes are often small in this literature and many findings have not been replicated. For these reasons, we conducted an empirical analysis, described below.

To fully identify and characterize the WM tract composition of each social brain pathway, we performed probabilistic tractography on a large DTI dataset (103 healthy young adult subjects). This dataset is based on our in-house DTI database, accumulated from previous studies, all using the same MR procedures and parameters (Alm, Rolheiser, Mohamed, & Olson, 2015; Alm, Rolheiser, & Olson, 2016; Hampton et al., 2016; Metoki et al., 2017; Unger et al., 2016). We choose probabilistic tractography because it enables us to estimate the likelihood/probability of every voxel involved in the trajectory of a defined WM pathway (Behrens, Berg, Jbabdi, Rushworth, & Woolrich, 2007). By overlaying this probabilistic map on a standard WM atlas (i.e. ICBM-DTI-81 atlas, Mori et al., 2008), we were able to extract the contribution of each known WM tract to each social brain pathway (see detailed methods description in Supplementary Materials and Methods). In short, our goal was to build the connectivity matrix between putative regions in each social brain network and elucidate the fiber tract composition for each pathway (see Figure 5 and Table 4-6).

**Figure 5.**
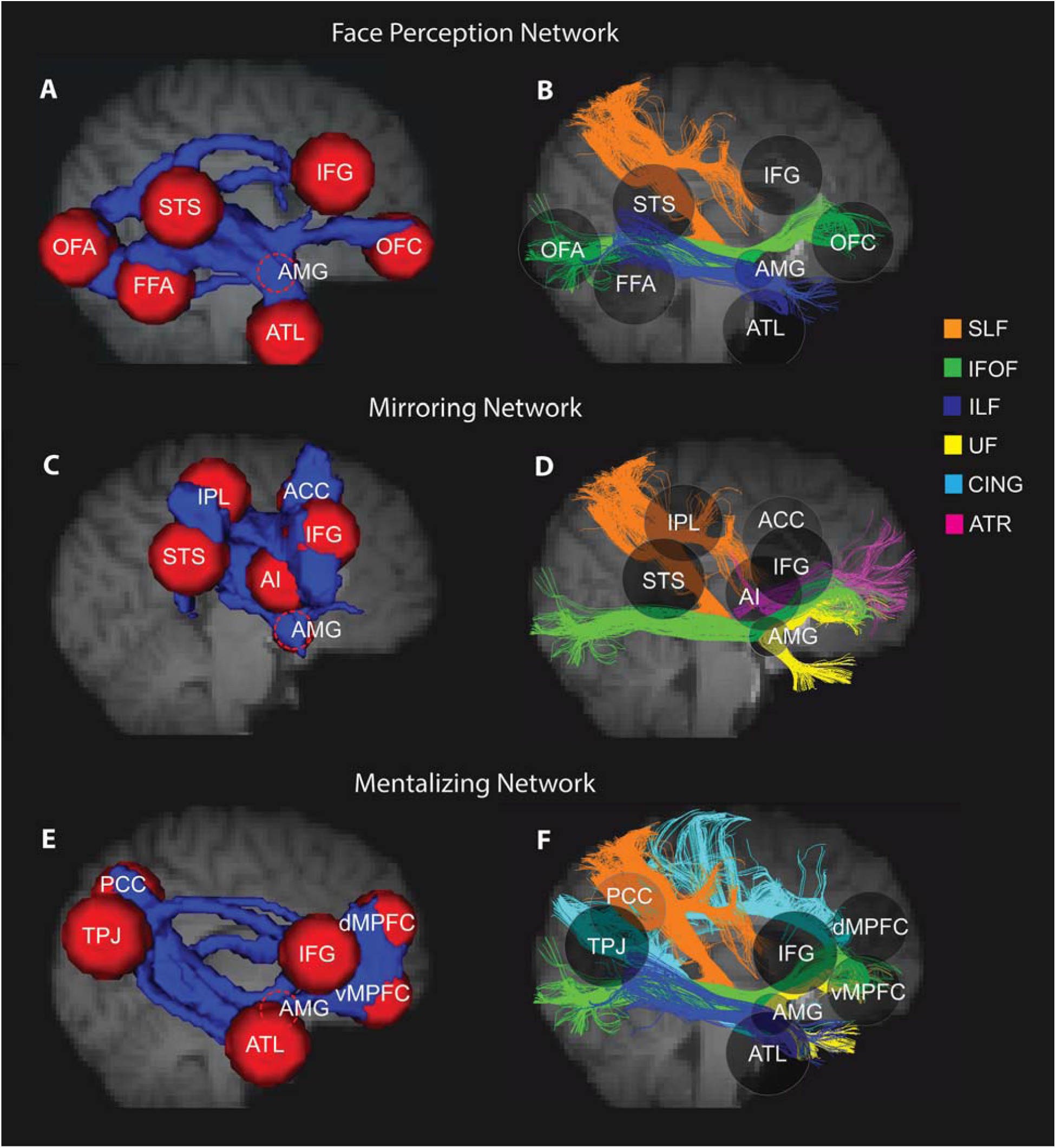
Social brain white matter tracts. Using probabilistic tractography, we reconstructed the WM skeleton, across 103 subjects, between putative regions in each social brain network (A)(C)(E), and we summarize the major white matter tracts for each network based on the literature review and the present tractography (B)(D)(F). In the left column, each red sphere represents a gray matter region of interest (ROI) and the blue represents the tractography-reconstructed WM pathways between ROIs. In the right column, transparent spheres are retained to use as landmarks. Different white matter tracts are represented by different colored streamlines.

**Table 4.**
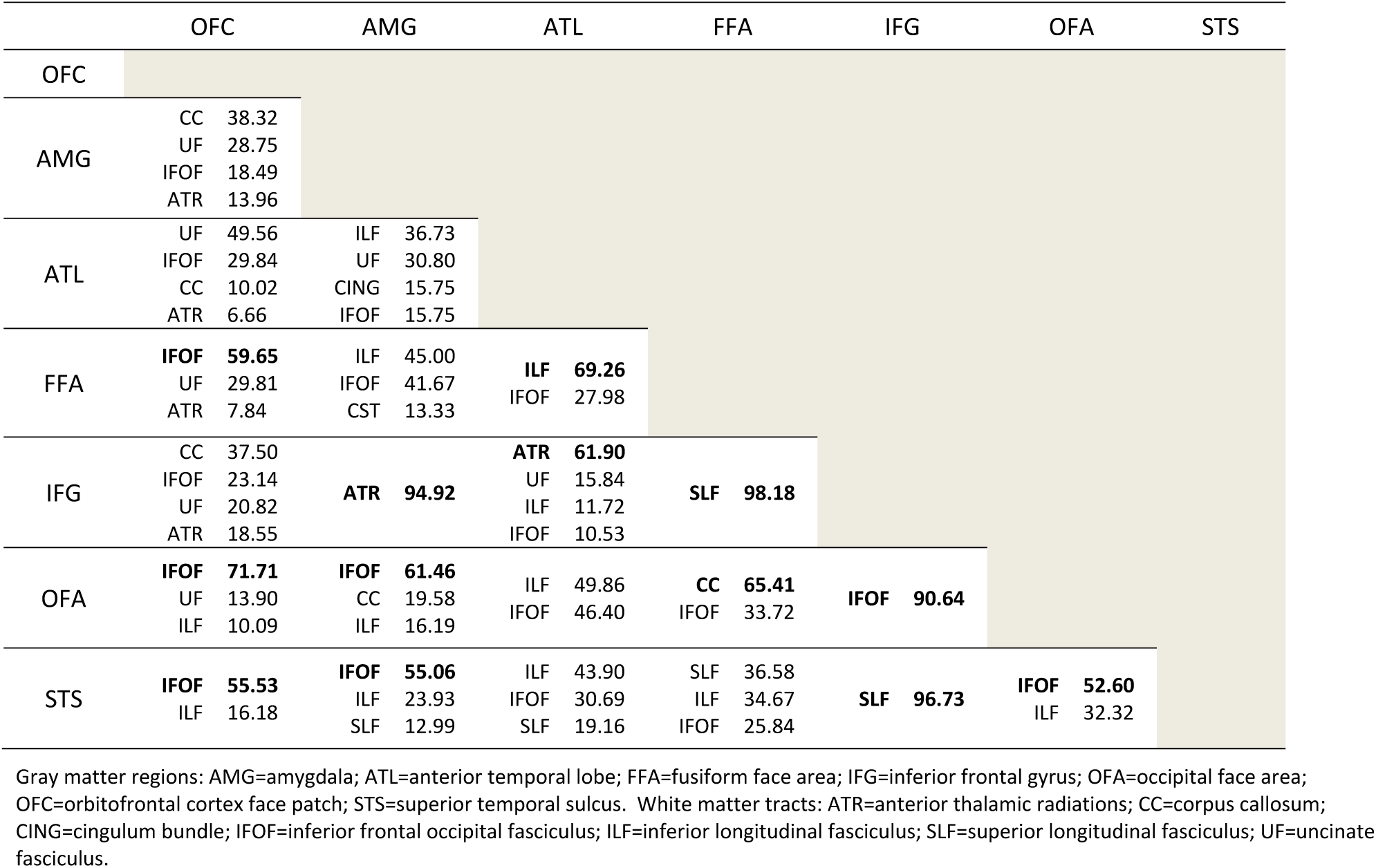
The connectivity matrix for the face perception network. All numbers are tract composition percentage (%). Only tracts with percentages larger than 5% are shown. Tract percentages greater than 50% are presented in bold font. The labels across the top and left side denote gray matter regions. White matter tracts are listed inside the connectivity matrix.

**Table 5.** The connectivity matrix for the mirroring network. All numbers are tract composition percentage (%).Only tracts with percentages larger than 5% are shown. Tract percentages greater than 50% are presented in bold font. The labels across the top and left side denote gray matter regions. White matter tracts are listed inside the connectivity matrix.

|  | ACC |  | AI |  | AMG |  | IFG |  | IPL |  | STS |
| --- | --- | --- | --- | --- | --- | --- | --- | --- | --- | --- | --- |
| ACC |  |  |  |  |  |  |  |  |  |  |  |
| AI | SLF | **86.20** |  |  |  |  |  |  |  |  |  |
|  | IFOF | 11.06 |  |  |  |  |  |  |  |  |  |
| AMG |  |  | SLF | 44.19 |  |  |  |  |  |  |  |
|  | ATR | **81.25** | IFOF | 28.42 |  |  |  |  |  |  |  |
|  | ILF | 7.55 | CST | 14.63 |  |  |  |  |  |  |  |
|  |  |  | UF | 10.85 |  |  |  |  |  |  |  |
| IFG | SLF | **96.10** | SLF | **94.75** | SLF | **85.06** |  |  |  |  |  |
|  |  |  |  |  | ATR | 8.79 |  |  |  |  |  |
| IPL | SLF | **99.57** | SLF | **84.87** | SLF | **75.54** | SLF | **99.54** |  |  |  |
|  |  |  | CST | 8.04 | CST | 22.21 |  |  |  |  |  |
| STS | SLF | **99.00** | SLF | **83.91** | SLF | **81.45** | SLF | **99.68** | SLF | **96.35** |  |
|  |  |  | IFOF | 11.63 | IFOF | 7.00 |  |  |  |  |  |
|  |  |  |  |  | CST | 6.73 |  |  |  |  |  |

Gray matter regions: ACC=anterior cingulate cortex; AI=anterior insula; AMG=amygdala; IFG=inferior frontal gyrus; IPL=inferior parietal lobe; STS=superior temporal sulcus. White matter tracts: ATR=anterior thalamic radiations; CST=cortico-spinal tract; IFOF=inferior frontal occipital fasciculus; SLF=superior longitudinal fasciculus; UF=uncinate fasciculus.

**Table 6.**
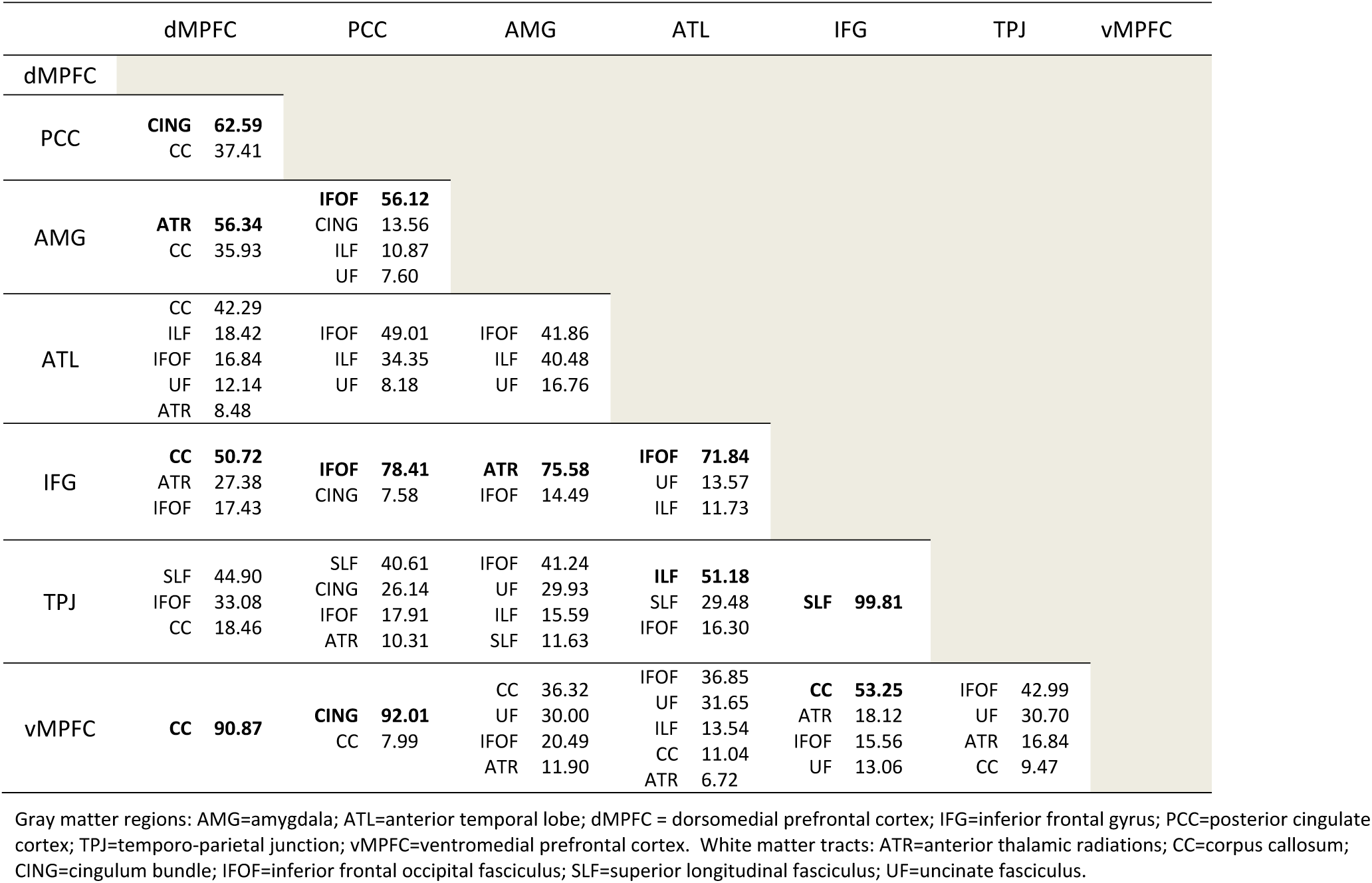
The connectivity matrix for the mentalizing network. All numbers are tract composition percentage (%). Only tracts with percentages larger than 5% are shown. Tract percentages greater than 50% are presented in bold font. The labels across the top and left side denote gray matter regions. White matter tracts are listed inside the connectivity matrix.

### Results

For the face perception network, probabilistic tractography revealed that 30.43% of WM voxels can be classified by the tracts listed in the ICBM-DTI-81 atlas. Among all classified tracts, the SLF occupied the most WM volume in the face network (*32.35%*), followed by the IFOF (*27.32%*), ILF (*23.81%*), CC (*6.17%*), ATR (*5.56%*), and UF (*5.35%*). When we more closely examined which specific pathways these tracts mainly subserved (see Table 4), we found that the SLF constituted a large proportion of two dorsal pathways projecting to the IFG (*FFA-IFG: 98.18%; STS-IFG: 96.73%*). This means that 98.18% of the voxels in the FFA-IFG pathway were classified as SLF, so were 96.73% of the voxels in the STS-IFG pathways. The IFOF was observed to mediate communications between posterior core face areas and anterior fronto-amygdala areas (*OFA-IFG: 90.64%; OFA-OFC: 71.71%; OFA-AMG: 61.46%; FFA-OFC: 59.65%; STS-OFC: 55.53%; STS-AMG: 55.06%*), and the ILF was found to support pathways along the ventral stream (*FFA-ATL: 69.26%; OFA-FFA: 65.41%; OFA-ATL: 49.86%*). In addition, the CC appeared to take part in two pathways with the OFC (*AMG-OFC: 38.32%; IFG-OFC: 37.50%*), and the ATR subserved connections between medial temporal cortex and IFG (AMG-IFG: 94.92%; ATL-IFG: 61.90%). Finally, the UF was found to be involved in connections between ATL, AMG and OFC (*ATL-OFC: 49.56%; ATL-AMG: 30.80%; AMG-OFC: 28.75%*)

For the mirroring network, only 16.46% of the WM voxels could be classified by the tracts listed in the ICBM-DTI-81 atlas. Among them, the SLF was the most dominant tract occupying 83.86% of WM voxels in the mirroring network, with the rest labelled as the IFOF (*6.72%*), corticospinal tract (CST, *5.17%*), ATR (*1.96%*), and UF (*1.47%*). For the tract composition of each pathway in the mirroring network, Table 5 shows that the SLF mediated all pathways between perisylvian regions (*STS-IFG: 99.68%; IPL-IFG: 99.54%; STS-IPL: 96.35%)* and played an important role in most ACC connections (*IPL-ACC: 99.57%; STS-ACC: 99.00%; IFG-ACC: 96.10%*). Albeit in smaller proportions, the IFOF was found to be part of AI-related pathways (*AMG-AI: 28.42%; STS-AI: 11.63%; AI-ACC: 11.06%*), and the CST was part of AMG-related pathways (*IPL-AMG: 22.21%; AMG-AI: 14.63%*). The ATR and UF were mainly involved in AMG-ACC (*81.25%*) and AMG-AI pathway (*10.85%*), respectively.

Last, for the mentalizing network, probabilistic tractography revealed that 32.04% of WM voxels were atlas-listed tracts. The CC had the largest proportion of volume (*32.58%*), followed by the IFOF (*18.80%*), SLF (*16.39%*), ATR (*14.93%*), ILF (*7.14%*), and UF (*5.79%*). As shown in Table 6, large percentages of voxels in major frontal pathways were labeled as the CC (*vMPFC-dMPFC: 90.87%, vMPFC-IFG: 53.25%; IFG-dMPFC: 50.72%*). The IFOF mediated several projections to the PCC (*PCC-IFG: 78.41%; PCC-AMG: 56.12%; PCC-ATL: 49.01%*), as well as to the ATL (*ATL-IFG: 71.84%; ATL-AMG: 41.86%*). The ILF was also part of ATL-related pathways (*TPJ-ATL: 51.18%; ATL-AMG: 40.48%; ATL-PCC: 34.35%*). The SLF exclusively subserved the connection between the TPJ and IFG (*99.81%*), and the ATR was the main tract for amygdala-frontal connections (*AMG-IFG: 75.58%; AMG-dMPFC: 56.34%*). In addition, the UF was engaged in vMPFC connections with the ATL (*31.65%*), TPJ (*30.70%*), and AMG (*30.00%*). Finally, although the cingulum (CING) occupied a small percentage of WM in the network (*4.33%*), it is the most dominant tract connecting all distant medial mentalizing areas (*PCC-vMPFC: 92.01%; PCC-dMPFC: 62.59%*).

These results (see a summary in Figure 5) are in line with previous studies using similar tractography methods. For the face perception network, there is a dorsal and a ventral pathway. The dorsal pathway runs from face-selective STS to the IFG via portions of the SLF (Ethofer et al., 2013; Gschwind et al., 2012). The ventral pathway runs from OFA and FFA to the ATL and AMG via the ILF and IFOF, and extends even more anteriorly into face-selective frontal areas (IFG and OFC) via the IFOF (Gschwind et al., 2012; Pyles et al., 2013). For the mirroring network, Hecht et al., (2013) and Hamzei et al., (2016) revealed that the SLF mediates two main pathways linking core mirroring areas (e.g. IPL-IFG, STS-IFG), and the ILF is involved in the STS-IPL connection. No DTI study thus far has directly elucidated the WM architecture of the mentalizing network; however, since the mentalizing network shares much overlap with the default mode network (DMN) both anatomically and functionally (Buckner, Andrews-Hanna, & Schacter, 2008; Li, Mai, & Liu, 2014; Mars, Neubert, et al., 2012), we can glean insights from that line of research. Most DMN studies show that the dorsal cingulum mediates the MPFC-PCC pathway, while the ventral cingulum supports communications between the PCC and ATL/AMG (Greicius, Supekar, Menon, & Dougherty, 2009; Sethi et al., 2015; van den Heuvel, Mandl, Kahn, & Hulshoff Pol, 2009; van den Heuvel et al., 2008). The TPJ connects with the ATL/AMG via the ILF (Horn, Ostwald, Reisert, & Blankenburg, 2014), and with the MPFC ventrally via the IFOF and dorsally via the SLF (Grosse Wiesmann et al., 2017; van den Heuvel et al., 2009). In the present study, we observed these reported structural connections and beyond that, we identified the whole connectivity profile of each social brain network, as well as the tract composition of each social pathway. In addition, our tractography results converge well with our literature review. The ILF, IFOF, and SLF are the most reported tracts for face processing in the literature, and in our tractography analyses they are the top 3 tracts occupying the majority of WM voxels in the face network. Past studies suggest that the SLF is the most important tract for embodied cognition, and our tractography results also support its dominant role in the mirroring network (i.e. 83.86% of WM voxels belong to the SLF). Finally, for the mentalizing network, the same tracts (i.e. cingulum, SLF, UF, IFOF, and ILF) were found in both the ToM literature and the current analyses.

We also encountered some unexpected findings, most of which related to the corpus callosum (CC). For example, the connectivity matrix of the face perception network (Table 4) revealed that the AMG-OFC pathway was mainly subserved by the CC (38.32%), followed by the UF (28.75%). Since all ROIs defined in our tractography were in the right hemisphere, this finding conflicts with our knowledge of brain anatomy, since the right AMG and OFC should be primarily connected by the right UF, rather than the CC (Catani & Thiebaut de Schotten, 2008). Similar erroneous findings can also be manifested in MPFC-related mentalizing pathways (Table 6), such as spuriously high probability of CC involvement in the dMPFC-vMPFC connection (90.87%). These problems might arise from our atlas-based approach, which computes the tract composition of each pathway by overlaying the probabilistic map onto a standard WM atlas. This method typically works well for WM connections with simple fiber configurations but is prone to produce artefactual results when the pathway travels through structures with high uncertainty of fiber orientations (i.e. crossing fiber sites), as is seen in the CC. Moreover, it is important to bear in mind that the tract percentage numbers in the connectivity matrices only reflects the relative contribution of each atlas-listed tract for the pathway; they could completely change from one atlas to another. The ICBM-DTI-81 atlas we used for the current analysis includes 48 tracts (Mori et al., 2008), which means we can only elucidate the tract composition based on these tracts. This is why the connectivity matrix of the mirroring network in Table 5 shows the sole engagement of the SLF for the STS-IFG and STS-IPL pathways although the literature suggests considerable involvement of the extreme/external capsule and the middle longitudinal fasciculus for these two pathways (Hecht et al., 2013). Since the ICBM-DTI-81 is currently the best atlas in use, future research should focus on developing more fine-grained WM atlases that include more segmented tracts and labels. This will profoundly increase the value and accuracy of atlas-based methods (including voxel-based analysis and TBSS).

## Current Problems and Challenges in the Field

Our review highlights the fact that there are already a large number of WM studies in the literature that enrich our understanding of the biological basis of social cognition. However, the methods and claims in many studies are far from perfect, which is to be expected, since diffusion imaging methods are continually improving. Here, we briefly describe some of the most common problems in the field, and offer some practical suggestions to help address these challenges. We find that limitations of experimental design, data acquisition, diffusion modeling, replicability, and results interpretation and reporting may underlie many of the reported effects. These limitations have the potential to undermine our understanding about the specific role of WM in social cognition, social behavior, and social disorders.

### Experimental Design

Choosing the proper behavioral paradigm to accurately and precisely measure the targeted social process is perhaps the biggest challenge in experimental design. Different social processes are inherently intertwined, thus most behavioral measures tap multiple constructs, making sub-processes difficult to disentangle (Happé, Cook, & Bird, 2017). For example, beside the embodied simulation process, facial emotion recognition tasks involve several other social processes including face discrimination, face classification, and the retrieval of emotion-related conceptual knowledge (Adolphs, 2016). Similarly, the comic strip vignettes task requires not only low-level action understanding abilities, but also high-level mental states attribution skills (Kana et al., 2014). It is problematic that most studies in the literature used only one task for their targeted social process and each used different tasks to probe the same alleged social process (see Tables 1-3). This makes it difficult to draw general conclusions across studies since different tasks might engage different nodes of the social brain network and thus implicate different WM connections (Grosse Wiesmann et al., 2017). For example, when measuring mentalizing functions, face-based ToM tasks, such as “reading the mind in the eyes”, evoke activations mostly in the IFG, while the story-based ToM tasks mostly recruit the MPFC (Schurz et al., 2014). When measuring empathy and facial emotion recognition abilities, the ACC engages mostly for pain scenes, whereas the insula and amygdala are recruited mostly for disgust and fear stimuli respectively (Bastiaansen et al., 2009). In the future, researchers should employ multiple measurements per study, such as using different stimuli (e.g. cartoons or stories), modalities (e.g. visual or verbal), paradigms (e.g. implicit or explicit ToM, Herbet et al., 2014; affective or cognitive empathy, Parkinson & Wheatley, 2014), task difficulty levels (Thomas et al., 2008), and methods (e.g. subjective self-reports/questionnaires or objective tools, such as using skin conductance responses to quantify empathic responses) when probing a targeted social process. This multi-measurement approach will help researchers make comparisons between studies and clarify the specificity versus generality of the role of a WM tract in a particular social process.

Another common drawback of experimental design in the literature is the lack of sufficient control variables measured in addition to the WM measurements. WM measurements can be measureably affected by participant’s intelligence (Penke et al., 2012), handedness (McKay, Iwabuchi, Häberling, Corballis, & Kirk, 2017), socioeconomic status (Ursache & Noble, 2016), and head motion (Yendiki, Koldewyn, Kakunoori, Kanwisher, & Fischl, 2014). Recent studies also show that WM structures underlying social cognition vary across age (Cabinio et al., 2015; Charlton et al., 2009; Thomas et al., 2008) and gender (Chou et al., 2011). These confounding factors should be explicitly matched between groups or regressed out during analysis. In addition, in order to examine the specificity of the relationships between the tracts of interest and target social processes, control tasks (e.g. non-social tasks with equivalent cognitive demands) and control tracts (Anderson et al., 2015; Unger et al., 2016) should also be included in a study’s design and analysis.

### Data Acquisition and Diffusion Modeling

Poor data quality in dMRI (e.g. noise, artifacts, and data under-sampling) often lead to errors in tensor estimation and, consequently, in diffusion maps that give rise to fiber reconstructions with erroneous orientations or lengths (Soares et al., 2013). Therefore, it is essential for dMRI researchers to acquire robust data with sufficient measurement and statistical power. Although imaging protocol optimization may vary based on the research question of interest, scanner hardware, scan time available, and specific anatomical structures of interest, here we provide some general guidelines for best practices in data acquisition. For example, it is recommended to sample along at least 30 unique gradient directions (even better, along 60+ directions) and to use a b-value of at least 1000 s/mm^2^ (even better, multiple b-values with some up to 2000–3000 s/mm ^2^) (Jones et al., 2013). In our literature review, more than 1/3 of the studies used protocols that did not meet these standards (Figure 4), which significantly undermines their reliability for obtaining robust estimates of tensor-derived properties or accurate reconstructions of WM tracts. We recognize that most of these studies examined developmental or clinical populations whose participants may have had difficulty withstanding longer acquisition times while being still (Scherf et al., 2014). For that, parallel imaging is one established way to reconcile these acquisition requirements while maintaining an acceptable acquisition time (Jones et al., 2013). Such simultaneous multi-slice (or multi-band) acquisitions drastically shorten measurement time and have been key innovations for improving data quality in recent large developmental diffusion imaging projects (e.g. the Developing Human Connectome Project, Hughes et al., 2016).

Another challenge for dMRI researchers is the wide variety of methods available to analyze the data. One must be wary of technical limitations and pitfalls when using a particular method to model WM patterns, especially in tractography (Jbabdi et al., 2015; Jones, 2010a, 2010b; Jones & Cercignani, 2010; Jones et al., 2013; Lerch et al., 2017). Human WM tissue is extremely difficult to model, due to its high density (e.g. more than 10,000 pathways with distinct origins and terminations, Jbabdi et al., 2015), complex trajectory patterns (e.g. 90% of WM voxels contains crossing fibers, Jeurissen, Leemans, Tournier, Jones, & Sijbers, 2013), and intricate organizations in its cortical origins/terminations (e.g. superficial WM bundles running parallel to the cortex, impeding the detection of fibers entering the cortex, Reveley et al., 2015). Despite these complexities, 50% of tractography studies in our literature review used a single tensor model with a deterministic tracing algorithm, which only calculated a single principle diffusion direction in each voxel. This method is too simplistic and inadequate for characterizing WM microstructure in crossing fiber voxels throughout the brain and can lead to erroneous fiber trajectory reconstructions. Optimal approaches include high-angular-resolution diffusion imaging (HARDI) with more complex biophysical models, such as multi-tensor models and ball-and-stick models, or utilizing advanced multi-shell “model-free” techniques such as Q-Ball Imaging and diffusion spectrum imaging (Wandell, 2016). These sophisticated tools are increasing popularity in other fields of neuroscience such as vision research (e.g. Rokem et al., 2017), but thus far, were only employed by three social neuroscience studies (Anderson et al., 2015; Olszewski et al., 2017; Pyles et al., 2013). We recommend that future research use these advanced methodologies whenever possible in diffusion data analysis, because they can significantly increase the sensitivity and fidelity of tractography (Jones et al., 2013), and most importantly, improve reliability and reproducibility.

Taken together, the evolution of our understanding of the WM pathways underlying social cognition depends intimately on the development of diffusion imaging techniques. As data acquisition, image processing, and diffusion modeling approaches continue to develop, our ability to identify the link between social processes and WM tracts will continue to improve.

### Replicability

In recent years, social psychology and neuroscience have been heavily criticized for their low replication rates (Open Science Collaboration, 2015; Vul, Harris, Winkielman, & Pashler, 2009). As a subfield, discrepancies in findings are not uncommon, particularly in dMRI studies. Since this technique provides indirect measurements of WM tissue and relies heavily on data quality and analysis methods, it can yield inconclusive and sometimes contradictory results. For example, Thomas et al., (2009) used DTI to evaluate the structural integrity of long-range visual tracts (i.e. ILF and IFOF) in individuals with developmental prosopagnosia (DP). They found that these patients showed reductions in the integrity of the right ILF and these reductions were positively correlated with individual face perception deficits. In contrast, Gomez et al., (2015) did not detect any WM integrity reductions in the right ILF in DP subjects; instead, they found that DP arises from the local WM difference in the right FFA, rather than any long-range WM tracts.

A few explanations can account for these discrepancies. One likely factor is the small sample size and concomitant low statistical power in both studies (n=6 in Thomas et al., 2009; n=8 in Gomez et al., 2015). Another possibility comes from the differences in data quality and analysis methods. Thomas et al., (2009) collected diffusion data with only 6 gradient directions and analyzed the data with simplistic deterministic tractography, while the more recent study by Gomez et al., (2015) employed 30 gradient directions and probabilistic tractography. These interpretations are further supported by a recent study (Song et al., 2015) using a larger sample size (n=16), optimized scanning parameters (61 directions), and multiple analysis methods (deterministic tractography, probabilistic tractograpphy, and voxel-wise comparison). Consistent with Gomez et al., (2015), they found no differences on any of the WM measures in the right ILF between the DP and control group, but local WM differences in the right FFA accounted for the face perception deficit in DP. Thus, it seems that small sample size, poor data acquisition, and the relatively simple tractography method prevented the findings in Thomas et al., (2009) from being replicated.

Moreover, we noticed that the literature used a variety of software packages to analyze dMRI data. It is important to note that not all software handles data in the same way (Christidi, Karavasilis, Samiotis, Bisdas, & Papanikolaou, 2016; Soares et al., 2013) and thus, can lead to differences in statistics and potentially the power to detect group difference and correlations between dMRI metrics and other measures (Jones et al., 2013). We believe that using well-documented open source software and making in-house algorithms transparent will help facilitate the comparison between different methods and studies and will ultimately, promote reproducible neuroscience.

In summary, as the sample size, data quality, and analysis methods are significantly improving, we expect replicability in this field to similarly improve (Poldrack et al., 2017). Two recent studies support our opinion by showing that one can get very similar connectivity profiles for the face perception network when utilizing advanced data acquisition and sophisticated tractography methods (Gschwind et al., 2012; Pyles et al., 2013). In addition, large open datasets are available for reproducible science. The Human Connectome Project (Van Essen et al., 2013) is an excellent example (e.g. > 1200 subjects already available) which provides not only high-quality dMRI data, but also social cognitive measurements from fMRI and behavioral tests. Future research should take advantage of this resource to provide more confident inferences about the role of WM in social cognition.

### Results Interpretation and Reporting

Finally, even when the researchers make the right choices for data acquisition and modeling, and the results are proven to be robust, reliable, and reproducible, we still need to improve the ways we interpret and report results in the literature. dMRI measures carry invaluable information about WM microstructure and are certainly sensitive indicators of WM changes and neuropathology. Many studies have observed FA differences between groups or correlations with social behavioral measures. However, the interpretation of such relationships is complex and should be performed with care. Although FA may reflect fiber integrity, it can also be confounded by factors that do not necessarily reflect WM integrity, such as partial volume effects (i.e., signal mixing of WM, gray matter, and cerebrospinal fluid), or heterogeneity in the orientation of axons (Jbabdi et al., 2015). In the absence of other information, FA is a fairly ambiguous and non-specific biomarker of WM microstructure and neuropathology (Alexander et al., 2007). Therefore, WM integrity assumptions based on FA must always be made with extreme caution. In addition, any notion of being able to relate FA to behavioral performance in a linear fashion is flawed, because higher FA is not always associated with superior social performance (Imfeld, Oechslin, Meyer, Loenneker, & Jancke, 2009; Unger et al., 2016). When exploring the WM correlates of individual differences, researchers should not misinterpret that those participants with higher FA have better WM connectivity, nor should they over-interpret that higher or lower FA is the cause of better performance on the tasks (Unger et al., 2016). Even in a situation in which higher FA indeed reflects superior WM connectivity, it is impossible to discern whether a pathway is inhibitory or excitatory. Therefore, lower or higher FA values must be interpreted in the context of the known functions of a pathway and its connecting regions (Roberts et al., 2013).

With regard to results reporting, future studies need to provide more detailed WM anatomy information about their findings. Large tracts like the SLF are composed of distinct subsections (e.g. SLF I—SLF III and AF, Kamali et al., 2014) and for each subsection, many fasciculi enter and exit at different points such that not all WM bundles traverse the full length of the tract. Since different subsections of a tract might be responsible for different cognitive functions (Metoki et al., 2017; Tavor et al., 2014), it is important for researchers to report sufficient information about the precise location of their results (e.g. anterior or posterior portion of the ILF; genu or splenium of the CC).

In addition, most studies in the literature only report results based on FA (Figure 4), which may not be enough to fully characterize the underlying changes in WM microstructure. Additional DTI measures, including MD, AD, RD, and volume, may be more informative regarding the specific nature of WM changes and dysfunction. It has been argued that all DTI measures should be routinely analyzed and reported even if some are not statistically significant (Alexander et al., 2007). We strongly support this practice and believe reliable interpretation of dMRI research requires not only sophisticated analyses, but also a complete and comprehensive report of WM anatomy and measures.

## Future Directions

In the next section, we are going to discuss five promising future directions. The first three are recommendations: that theories should be tested, that WM should be examined at a finer scale, and that dMRI can be combined with other imaging methods to improve inferential power. The last two are topical areas that deserve more focused research: the importance of local WM and cerebellar WM for social cognition.

### Testing Current Theories in Social Neuroscience

Research on WM is opening new avenues to test theories in social neuroscience. Theories that are most amenable to this endeavor are ones that propose some sort of ordered processing. For instance, the Haxby model is considered the most dominant neural theory of face processing (Gobbini & Haxby, 2007; Haxby et al., 2000). According to this model, the OFA, FFA, and STS constitute the core system that subserves the visual analysis of faces, and more anterior areas, such as the ATL, AMG, and OFC comprise the extended system that gleans other information from faces, such as their emotional and personal significance. This model postulates a hierarchical structure such that the OFA projects to both the FFA and STS, and each plays a different role in face processing (Bernstein & Yovel, 2015). After being processed by the core system, information is then sent to the extended system to extract biographical knowledge, analyze emotional information, or to evaluate facial attractiveness.

However, recent WM research challenges this framework and suggests the need for modifications to this model (Duchaine & Yovel, 2015). First, there is evidence that the OFA is not the only entry point for the face processing network. A DTI study using probabilistic tractography (Gschwind et al., 2012) revealed several direct WM pathways between early visual cortices and face-selective regions without any mediation by the OFA, and some of these pathways were even stronger than connections with the OFA. This signifies that face processing might not proceed in sequence, but rather in a parallel and interactive fashion. Second, there is evidence for multiple WM pathways to each face-selective region, via distinct sub-tracts of the ILF or IFOF. This helps explain cases of neuropsychological patients who have lost bilateral OFA but FFA and STS remain sensitive to faces (Duchaine & Yovel, 2015), as well as patients who have lost most of the core face-processing system but exhibit preserved activations in the extended system (Valdés-Sosa et al., 2011). Converging evidence for this new view also comes from magnetoencephalography research which has suggested that some “short-cuts” for rapid communication exist between early visual cortices and more anterior face areas involved in affective processing (e.g. AMG and OFC), allowing attention to be modulated before the face stimulus has been fully processed by the ventral stream (Barbeau et al., 2008; Rudrauf et al., 2008).

Additionally, the Haxby model argues for dynamic interactions between regions in the core system, particularly with the OFA providing inputs to both the FFA and the STS. However, previous studies reported strong correlations between the time courses of OFA and FFA activation but much weaker correlations between activation of these areas and the STS (Avidan et al., 2014). TMS studies also found that stimulation to the OFA influenced the neural response of the FFA but not the STS (Pitcher, Duchaine, & Walsh, 2014). These observations imply that the OFA and FFA are tightly connected, but the STS seems to be more isolated within the core system. Two dMRI tractography studies confirmed this claim by demostrating that direct WM connections only exist between the OFA and FFA, but not between the OFA and STS or between the FFA and STS (Gschwind et al., 2012; Pyles et al., 2013). They also found that the STS is preferentially connected to the SLF fiber network rather than to the ILF/IFOF network by which the majority of face processing areas are connected. Together these findings warrant a revised framework to the Haxby model: in the new model, the OFA and FFA constitute part of a ventral pathway of face processing, while the STS is part of a dorsal pathway originating from distinct early visual areas involved in motion processing (Bernstein & Yovel, 2015; Duchaine & Yovel, 2015; Pyles et al., 2013).

As we can see, WM investigation provides a unique way to gain insights into the underlying organization of the face perception network, beyond the traditional mapping approaches based on functional activations alone. This approach has only been applied to a small number of theories, such as the dual-stream model of empathy (Herbet, Lafargue, Moritz-Gasser, Menjot de Champfleur, et al., 2015; Parkinson & Wheatley, 2014) and mentalizing (Herbet et al., 2014). Future research should continue to test theoretical models in social neuroscience from the perspective of structural connectivity.

### Examining WM at a Finer Scale

Another important future direction will be to investigate the functional segmentation of particular WM tracts. As we noted earlier, the ILF has been implicated in face processing, empathy, emotion recognition, and mentalizing abilities. This seemingly nonspecific role of the ILF in a variety of social processes may not be surprising, considering that the ILF is a large fasciculus reaching up to 12cm in length and that different fiber bundles enter and exit the fasciculus at various positions. As such, the properties of WM tissue vary systematically along the trajectory of the ILF, potentially yielding distinct functional subcomponents that support discrete social functions. For example, Tavor et al., (2014) reported the anterior portion of the ILF is associated with face memory abilities, whereas the middle and posterior portions are associated with scene memory abilities. Gomez et al., (2015) revealed that WM properties near the face-selective regions (i.e. FFA) are specifically correlated with an individual’s face recognition ability, while WM properties near place-selective regions are correlated with place recognition ability. In addition, Song et al., (2015) identified that the posterior portion of local FFA fibers (and not the anterior portion) is associated with the face recognition deficits in prosopagnosia patients. These findings all suggest the existence of segregated segments or pathways within the ILF, each specialized for distinct functions (e.g. face vs. scene processing).

In most studies, WM measurements (e.g. FA, MD) were extracted and analyzed across the entire tract. Such large-scale approach is crude and limits our ability to examine how WM relates to functional regions or relays information specific to a social process. More recently, researchers have begun to examine WM properties from different positions or segments within a tract to reveal variations within a specific tract (Yeatman, Dougherty, Myall, Wandell, & Feldman, 2012). This approach of quantifying WM properties within each component of a fiber tract may provide new insights into WM function or dysfunction that are not obvious from the mean measures of that tract, and thus has the potential to reveal a great deal about the functional organization of fibers in the social brain. We contend that future studies in social neuroscience should explore the fine-grained functional segmentation of the WM tracts identified in this review (e.g. SLF, cingulum, UF). Instead of using traditional anatomical approaches to define entire fasciculi (Catani & Thiebaut de Schotten, 2008), researchers should consider utilizing functional ROIs as seeds in tractography to identify the sub-tracts of large fasciculi that are specific to particular social processes (e.g. the pathway between the face-selective STS and IFG for social gaze perception, Ethofer et al., 2011). These functionally defined WM tracts will provide us a more detailed and refined picture of the structural architecture for social cognition.

### Collecting Multimodal Brain Data to Study Social Cognition

Collecting multimodal brain data from the same individual using different neuroimaging methods has recently become a standard in human neuroscience research and is certainly a trend for the future (Soares et al., 2013). Combining dMRI with other modalities allows for a comprehensive view of human brain structure and function and has promising clinical applications (Meng et al., 2017). For example, dMRI can be combined with advanced sMRI techniques (e.g. magnetization transfer, susceptibility-weighted imaging, and quantitative MRI) to reveal more fine-grained features of WM microstructure, such as axon density, diameter, and myelin sheath thickness (Lerch et al., 2017; Wandell, 2016). This helps to form a better and more complete assessment of social WM structures and provides invaluable insights about how social brain WM tracts are altered in development, aging, and disorder. Similarly, dMRI and fMRI can be complementarily integrated to gain a more precise understanding of functional brain networks. We can use functional connectivity to infer information regarding anatomical connections (Deco et al., 2014) or use structural connectivity to guide the construction of neurobiologically realistic models of functional connectivity during computational modeling (Stephan, Penny, Daunizeau, Moran, & Friston, 2009). Since functional connectivity is not affected by complex trajectory patterns (e.g. crossing fibers) or algorithm parameters (e.g. curvature and stopping criterion), it may allow us to indirectly validate the accuracy of tractography measures. Ultimately, we can explore the convergence of connectivity inference between tractography and functional connectivity to examine whether the same pattern of connectivity is confirmed by both modalities (Jbabdi et al., 2015).

Within our literature review, 17 out of 51 studies collected multimodal data (see Tables 1-3). Most of them implemented fMRI simply as a functional localizer to define social-specific ROIs for dMRI tractography. Three studies used DTI-fMRI to examine the relationship between structural and functional connectivity during social processing (Ethofer et al., 2011; Kana et al., 2014; Marstaller et al., 2016). One developmental study combined DTI with EEG (Taddei, Tettamanti, Zanoni, Cappa, & Battaglia, 2012) and revealed a longitudinal relationship between WM organization in adolescence and cerebral responses (N400) to hostile facial expressions in childhood. In line with facilitating such multimodal approaches, the Human Connectome Project provides notably high-quality dMRI and social-task-based fMRI and MEG data from a large number of healthy adults (Glasser et al., 2016). This excellent multimodal open resource should be exploited more by social neuroscientists. As more multimodal studies are conducted, we believe that convergent evidence from different techniques (e.g. dMRI, sMRI, fMRI, MEG) and fields (e.g. development, psychiatry, neurology) will grant us a global and complementary overview of the neurobiological basis of social cognition.

### Local Reginal WM for Social Cognition

It is important to note that the present literature has focused almost exclusively on large, major fasciculi but has ignored the contribution of local regional WM to social cognition. However, long-range WM tracts only comprise 4-10% of the whole human WM connectome, and the majority of WM consists of short association fibers in superficial WM (e.g. U-shaped fibers) that lie immediately beneath the gray matter and connect adjacent gyri (Jbabdi et al., 2015; Schuz & Braitenberg, 2002; Sotiropoulos & Zalesky, 2017; Wandell, 2016). Indeed, our probabilistic tractography results indicated that 60-80% of WM voxels in social brain networks are unclassified WM tissue, or in other words, do not belong to any atlas-listed long-range tracts. Technically, it is not easy to study local regional fibers, because their spatial arrangements around sulci or gyri are complicated and most dMRI tractography algorithms are unsuitable for reconstructing them (Feldman et al., 2010; Reveley et al., 2015). Researchers usually have to adopt voxel-based or ROI-based approaches to estimate local WM properties associated with social cognition. For example, Chou et al., (2011) used TBSS to examine local WM correlates of empathy ability and found that empathy quotient scores are positively correlated with FA in regional WM near social brain regions, such as the IPL and STS (see also Nakagawa et al., 2015; Takeuchi et al., 2013). Another way to accurately identify and characterize U-shaped fiber systems is to use novel fiber clustering algorithms (Zhang et al., 2014) or a sophisticated ensemble tractography approach (Takemura, Caiafa, Wandell, & Pestilli, 2016). For example, by using ensemble tractography to distinguish local and long-range WM streamlines, Gomez et al., (2015) revealed that properties of the local FFA WM, rather than properties of entire ILF, correlated with face-processing ability, and this relationship was abnormal in prosopagnosia patients. This finding has been replicated by another study using simple voxel-based analysis (Song et al., 2015). In the future, we anticipate more investigations of local WM function (especially those near social brain regions), as the present literature indicates such local WM can be critical to social processing in both healthy and clinical populations.

### Cerebellar WM for Social Cognition

Another important but unresolved issue concerns the role of cerebral-cerebellum structural connectivity in social cognition. Prior research suggests that the cerebellum is important for social cognition and certain types of cerebellar pathology can cause social deficits (Hoche, Guell, Sherman, Vangel, & Schmahmann, 2015; Van Overwalle, Baetens, Mariën, & Vandekerckhove, 2014, 2015; Van Overwalle, D’aes, & Mariën, 2015). However, our knowledge of the functional significance of cerebellar gray matter and WM is in its infancy. We have only limited knowledge of how social brain regions and the cerebellum are structurally connected (Sokolov, Erb, Grodd, & Pavlova, 2014) and how their structural connectivity is related to social function processing (Van Overwalle & Marien, 2016). Understanding the social WM pathways between cerebrum and cerebellum will help us clarify their functional relationship in social cognition.

## Limitations of the Current Review

We would be remiss if we didn’t point out some limitations of our review. First, we defined three social brain networks for paper classification. Although these networks are conceptually specialized for distinct social processes, they are not mutually exclusive. They overlap in regards to anatomy (e.g. AMG, IFG, STS/TPJ, see Figure 2) and interact with each other during many social tasks (Barrett & Satpute, 2013; Becchio et al., 2012; Greven & Ramsey, 2017; Sperduti, Guionnet, Fossati, & Nadel, 2014; Spunt, Satpute, & Lieberman, 2011; Zaki, Hennigan, Weber, & Ochsner, 2010). The way we classified each social task or process into three brain networks might be debatable, and there may be other ways to divide up social cognition. The challenge is that social processes are interdependent and multifaceted, and we are far from having an agreed-upon taxonomy or factor structure (Happé et al., 2017). For instance, some studies believe the “reading the mind in the eyes” task is measuring the mirroring network (Herbet, Lafargue, Moritz-Gasser, Bonnetblanc, et al., 2015), while others argue the task is probing the mentalizing network (Mike et al., 2013). Some researchers might categorize studies of “cognitive empathy”, a subtype of empathy, into the mirroring network group, while others would sort them into the mentalizing network because the term is conceptually interchangeable with perspective-taking and ToM (Shamay-Tsoory, 2011). Even when empathy as a whole belongs to embodied cognition, not all of its sub-components (e.g. “empathic concern”, “fantasy”, “personal distress”, see “interpersonal reactivity index”, Davis, 1983) are subserved by the same neural network (Kanske, Bockler, Trautwein, & Singer, 2015; Lamm & Majdandžić, 2015) or tracts (Fujino et al., 2014; Parkinson & Wheatley, 2014).

Second, we defined ROIs for our probabilistic tractography using mean MNI coordinates from prior meta-analysis studies. This analytic choice was imposed on us by using an existing dataset that lacked certain features. Ideally, functionally-defined gray matter regions should supplement dMRI, allowing for the creation of functional seeds for precise fiber tracking (Sotiropoulos & Zalesky, 2017).

Finally, several social processes were not covered by the current review due to the small number of studies in these domains examining WM indexes. Individual studies exist examining self-processing (Chavez & Heatherton, 2017), personality (Cohen et al., 2008), social reward (Bjornebekk, Westlye, Fjell, Grydeland, & Walhovd, 2012), peer influence (Kwon, Vorobyev, Moe, Parkkola, & Hamalainen, 2014), in-group bias (Baumgartner, Nash, Hill, & Knoch, 2015), social communication skills (Lo, Chen, Hsu, Tseng, & Gau, 2017), social decision-making (Barbey et al., 2014), and social network size (Hampton et al., 2016; Molesworth et al., 2014). Future reviews should include and discuss these in order to provide a larger overview of the WM basis for social cognition.

## Concluding Remarks

Research on structural connectivity in social neuroscience is a promising field for insights into the anatomical basis of social cognition, social behavior, and social disorders. In the present article, we comprehensively reviewed past literature to summarize the reported WM structures associated with social cognition and also empirically employed probabilistic tractography to elucidate major WM tracts scaffolding social brain networks. These two approaches demonstrated a converging group of tracts critical for face processing (the ILF, IFOF and SLF), embodied cognition (the SLF, UF, ATR and IFOF), and ToM (the cingulum, SLF/AF, UF, IFOF and ILF) (Figure 5). In addition, our review introduces multiple facets of WM research in social neuroscience, covering a wide array of approaches and applications. They all demonstrate the ways in which studying WM connectivity leads to a more complete understanding of the neurobiology of social cognition, its structure, functional properties, and relation to behavior and disorder. However, the main bottleneck of this exciting field is still the limited sample size, poor data quality, and simplistic analysis methods, which could be potentially addressed by utilizing large open datasets, such as the Human Connectome Project. Nevertheless, we are optimistic that the current paradigm shift towards connectivity will bring with it higher-quality data and a larger corpus of findings relevant to WM and social neuroscience.

